# Cellular state determines the multimodal signaling response of single cells

**DOI:** 10.1101/2019.12.18.880930

**Authors:** Bernhard A. Kramer, Lucas Pelkmans

## Abstract

Numerous fundamental biological processes require individual cells to correctly interpret and accurately respond to incoming cues. How intracellular signaling networks achieve the integration of complex information from various contexts remains unclear. Here we quantify epidermal growth factor-induced heterogeneous activation of multiple signaling proteins, as well as cellular state markers, in the same single cells across multiple spatial scales. We find that the acute response of each node in a signaling network is tightly coupled to the cellular state in a partially non-redundant manner. This generates a multimodal response that senses the diversity of cellular states better than any individual response alone and allows individual cells to accurately place growth factor concentration in the context of their cellular state. We propose that the non-redundant multimodal property of signaling networks in mammalian cells underlies specific and context-aware cellular decision making in a multicellular setting.

---

State-dependent decision making by individual cells is a hallmark of multicellular life. This is prominently exemplified in embryonic development, where spatial and temporal cues determine cellular fate, driving the emergence of complex multicellular structures which consist of multiple cell types with highly diversified patterns of gene expression^1–6^. Integration of spatial cues and cellular context is also observed *in vitro*. In intestinal organoids, symmetry breaking is determined by the mechanical context of individual cells^7^, while cell fates in pluripotent stem cell populations are determined by differences in cell density and distance to colony edge^8, 9^. Also in tissue culture cells, cellular activities and gene expression show predictable adaptation to heterogeneous microenvironments and cellular states^10–14^. The fidelity of these decisions relies on information processing by signaling networks^15–19^. This involves the concomitant activation of multiple connected signaling nodes composed of kinases, phosphatases, small GTPases, and effector proteins. The flow of information through these signaling nodes is highly variable between individual cells, even when cells are genetically identical and exposed to identical amounts of cytokine or growth factor^16, 20, 21^. As a result, cell populations can exhibit broad distributions of single-cell signaling responses^16, 22–25^. Although different concentrations of cytokine or growth factor indeed elicit distinct mean response values, an accurate measurement of the exact concentration is impossible for individual cells because the full response distributions overlap considerably^20, 25^ (**Fig. 1a**). It therefore remains largely unclear how individual cells are able to accurately integrate the complexity of variable extra- and intracellular information and convert this into a specific, context-aware response with the remarkable and reproducible accuracy necessary to sustain multicellular life.

**Fig 1.**
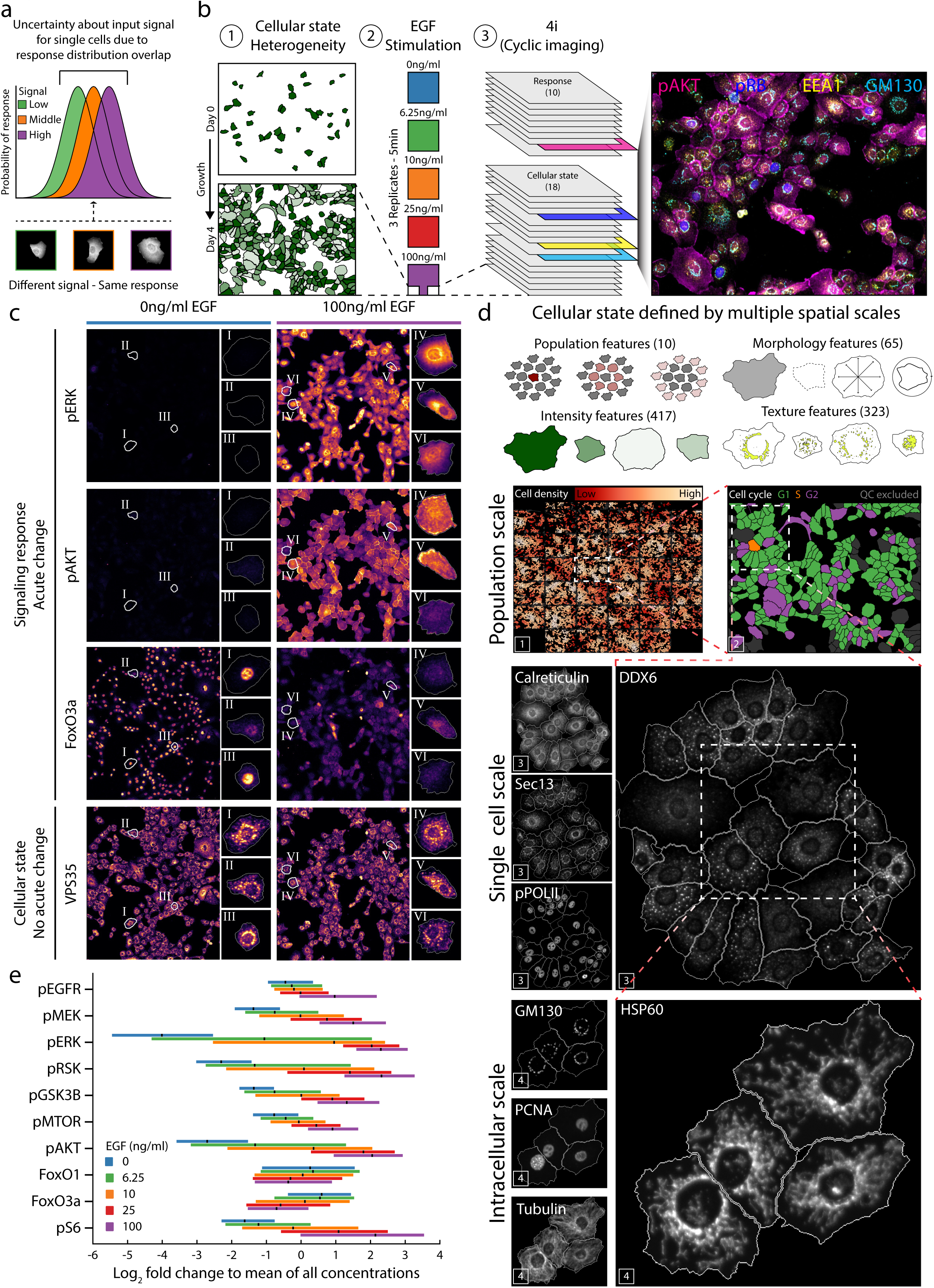
Capturing signaling responses and cellular state across spatial scales. (a) Schematic representation of overlapping response distributions. Populations of cells exhibit heterogenous signaling responses. The resulting distributions can display substantial overlap for signals of different strengths. This overlap seemingly impairs the ability of single cells to accurately estimate the signal strength. (b) Schematic representation of the experiment. Monoclonal, epithelial derived cells from mammary gland (184A1) were plated in low density in a 96-well plate and grown 4 days resulting in heterogeneity of local cell density. Cells were then depleted of serum and growth factor for 12 hours before exposure to 5 concentrations of EGF in 3 independent replicates for 5 min. 4i was performed with 18 antibodies against cellular state markers and 10 antibodies against response markers. The resulting 30-dimensional image (x/y + antibody stains) can be visualized as arbitrary combinations of those dimensions. One possible combination is displayed. (c) Visualization of signaling response and cellular state markers obtained by 4i in representative populations exposed to either 0ng/ml or 100ng/ml EGF for 5 min. Same populations are displayed for each marker and condition. Representative individual cells are highlighted (white segmentation outline, roman numerals) and are displayed next to population images. Same cells are displayed for each marker and condition. (d) 4i enables the characterization of a multivariate cellular state which is arises at multiple spatial scales. Assembly of individual imaging sites within one well (marked with a white 1) enables quantification of population scale features such as local cell density. Information about each cell’s cell cycle phase can be obtained using data from individual imaging sites (marked with a white 2). Heterogeneity of protein abundances become apparent at the single cell scale (marked with white 3s). Distinct subcellular localizations of proteins are revealed when assessing the intracellular scale (marked with white 4s). In combination, the features derived at these scales characterize each cell’s multivariate cellular state. (e) Single-cell percentile ranges (2.5 – 97.5) for each concentration of EGF and response marker. For each response marker, the mean across all concentrations of EGF and replicates was calculated. The ratio (fold change) of each cell to this mean was calculated. Percentile ranges of the log_2_ fold changes are then calculated for each condition (i.e. concentrations of EGF) and displayed as colored bar. Mean value of each condition is depicted in black.

One mechanism is based on temporal integration of the dynamic signaling response, which can provide more information than simple sampling of single time-points^21, 26, 27^. This, however, requires storage of information, as proposed for the plasma membrane or in the form of slowly accumulating effectors^28–30^, which may not facilitate an accurate response at short timescales. Another mechanism to accurately integrate complex inputs could rely on multimodal signaling and perception, a common principle in ecology and neuroscience. In multimodal perception, simultaneously incoming information of separate sensory modalities is integrated as a characteristic percept that is more comprehensive than each part alone^31, 32^. In these systems, activation of each modality occurs in a different context (e.g. the context of auditory and visual cues). Also in intracellular signaling, the context in which the activation of signaling nodes occurs is important^33, 34^. This is for instance seen in the phosphorylation of ERK on endosomes^35–37^ and its interaction with microtubules^38^, or in the modulation of MTOR activity by mitochondrial components^39, 40^, as well as in the effect of local cell density on the vesicular trafficking of EGFR^41^, and the activation of AKT on specific intracellular membranes^42, 43^. In each case, specific subcellular environments, which can vary depending on the multicellular context and cellular state^44^, are important in tuning signaling responses. Thus, the specific, non-redundant heterogeneity in the activation of individual signaling nodes of a network could provide an essential level of information which encodes cellular state and context characteristics. Collectively, as a multimodal response, these could generate a more accurate representation of the combined extra- and intracellular inputs than each signaling node alone.

## Capturing signaling responses and cellular state across spatial scales

Addressing this question requires quantitative information about the properties that comprise a specific cellular state, which must be obtained at subcellular and cellular scales, as well as at the scale of self-organized cell populations. It further requires the quantification of multiple signaling responses in a signaling network within single cells and how this response fluctuates across tens of thousands of cells. Measuring such a comprehensive amount of scale-crossing cellular features requires multiplexed protein measurements across multiple length units, recently enabled by 4i^44^, which we apply here to populations of monoclonal, human epithelial cells derived from mammary gland (184A1). 4i is unique in also achieving highly reproducible quantification at organellar resolution necessary to infer specific cell states^44^. Repeated measurements of phosphorylated ERK across multiple conditions demonstrated that 4i also achieves highly reproducible quantification of activated signaling proteins across large numbers of single cells (Pearson’s r > 0.98; Cells ∼3 x 10^5^; n = 15) (**Extended Data Fig. 1a**).

To address the question whether individual cells exhibit cell-state specific signaling responses which accurately correspond to variable growth factor inputs, we exposed 184A1 cells to 5 concentrations of epidermal growth factor (EGF) (0, 6.25, 10, 25, and 100 ng/ml) for 5 min, and performed 4i against 18 cellular state markers and 10 signaling response markers (**Fig. 1b; Extended Data Table 1,2**) in three independent replicates. The cellular state markers act as proxies of properties of the cytoskeletal and mechanical state of cells (Paxillin, Actin, Tubulin, Yap1), peroxisomes (ABCD3), endocytic (Dynamin, EEA1, VPS35) and exocytic (Calreticulin, Sec13, ERGIC53, GM130) organelles, position in the cell cycle and proliferative state (CyclinB, PCNA, pRB), transcriptional state (pPOLII), P-bodies (DDX6), and mitochondria (HSP60). The signaling response markers report on EGF-induced signaling, which include a phosphorylated form of the EGF receptor (pEGFR), 6 signaling kinases (pAKT, pGSK3B, pMEK, pERK, pMTOR, pRSK), the phosphorylated S6 subunit of the ribosome (pS6), and 2 transcription factors (FoxO3a and FoxO1) (**Fig. 1c,d; Extended Data Fig. 1b,c**). Each signal was quantified in ∼3x10^5^ individual cells using spinning disk confocal microscopy, followed by cell segmentation and feature extraction by means of computer vision and machine-learning (**Extended Data Fig. 1d**). A total of 815 features captured cellular properties apparent at different spatial scales, such as the single-cell abundances of each of the cellular state markers, their subcellular texture and shape properties, as well as properties of the multicellular scale, including local cell density and cell contacts. Collectively, we consider these a quantitative multivariate description of the cellular state (**Fig. 1d**).

The measurements revealed that response markers exhibited a log_2_ fold change in abundance of -1 to 6 upon EGF stimulation. As expected, abundance of the cellular state markers did not change upon stimulation and can therefore be used as a proxy for the cellular state before stimulation (**Extended Data Fig. 1e; Extended Data Table 3**). In addition, the response distributions exhibited an average overlap of ∼0.91 across replicates, illustrating high technical reproducibility (**Extended Data Fig. 1f**). Each response marker showed a typical dose-response relationship in their mean response, as well as extensive and varying heterogeneity in single-cell responses, with 2.5 – 97.5 percentile ranges covering a fold change of up to ∼80 (e.g. pERK at 6.25ng/ml EGF) (**Fig. 1e**). These heterogenous single-cell responses give rise to distributions that exhibit substantial overlap across distinct EGF concentrations, as previously observed (**Extended Data Fig. 1g**).

## Cellular state determines the heterogeneous signaling response

To test if heterogenous single-cell responses are determined by the cellular state, we first focused on the pERK response elicited by 6.25ng/ml EGF, as it exhibits the highest degree of heterogeneity (**Fig. 1e**). The collective of single cells displayed a broad distribution of pERK, including two distinct peaks (**Fig. 2a; Extended Data Fig. 1g**) as previously reported for low doses of EGF^23^. Using a Gaussian mixture model, we separated the cells into low- and high-responders. We then asked whether the observed qualitative difference in the acute response of pERK is linked to the pre-existing heterogeneity in cellular states in the population. Using logistic regression, we found that the propensity of individual cells to be a high or low responder could be accurately predicted based on their cellular state (F-score^Low-responder^: 0.95, F-score^High-responder^: 0.94) (**Fig. 2b; Extended Data Fig. 2a**). The features that allow this prediction arise at multiple spatial scales, such as local cell density and abundance of early endosomes (EEA1) (**Fig. 2c; Extended Data Fig. 2b**). Importantly, these features achieve an accurate prediction only when in combination, indicating the fundamental multivariate nature of cellular adaptation, which is invisible to univariate approaches (**Fig. 2d**). We similarly find that the two-peaked response distribution of pAKT (6.25ng/ml and 10ng/ml EGF) is well explained by properties of the cellular state. (**Extended Data Fig. 2c,d)**.

**Fig. 2.**
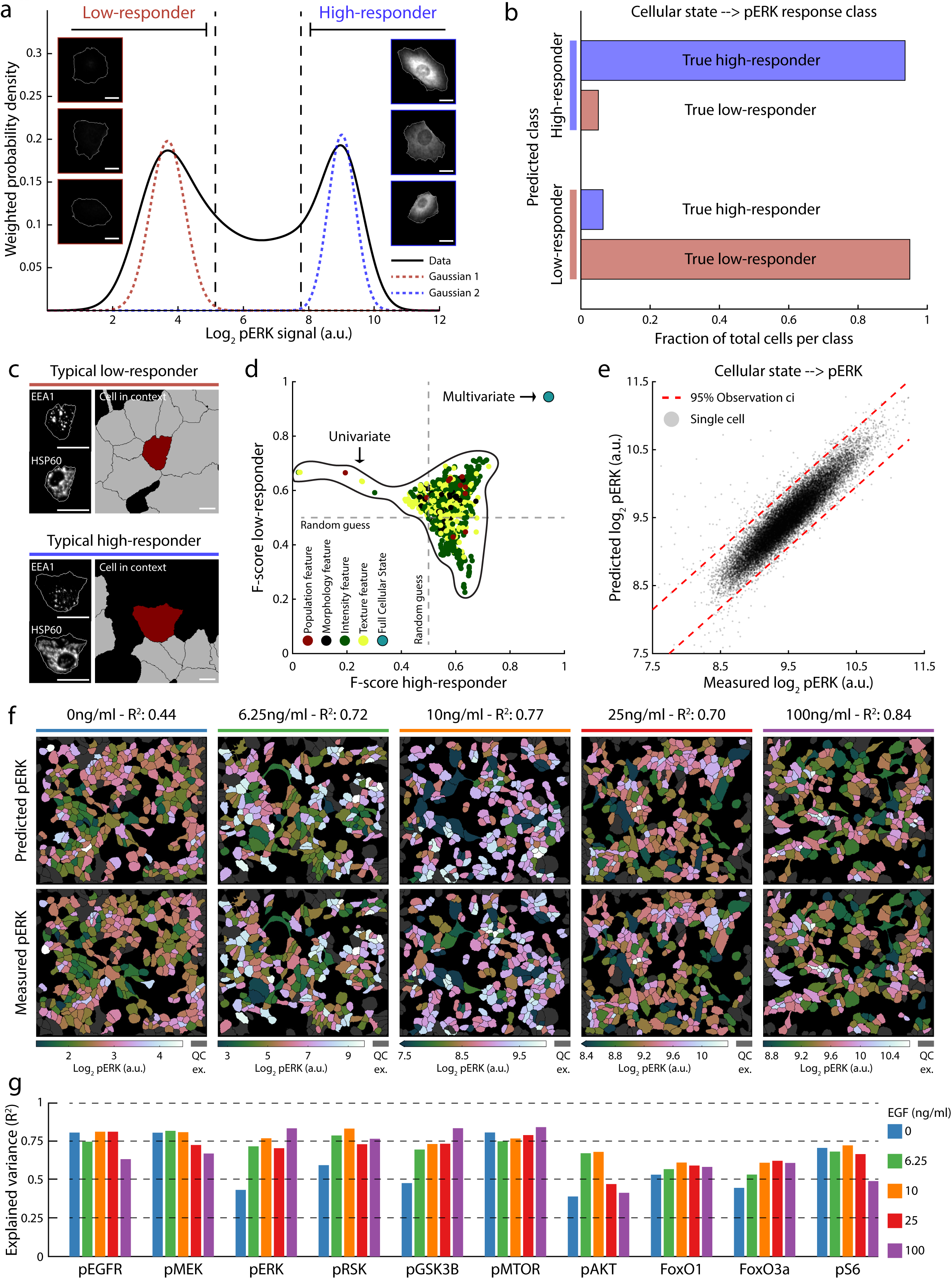
Cellular state determines the heterogeneous signaling response. (a) Phosphorylation of ERK in cell populations exposed to 6.25ng/ml of EGF exhibits two distinct peaks. Kernel density estimate with a gaussian kernel of the data is displayed. Cells were separated into low- and high-responder cells based on a gaussian mixture model (2 Gaussian components displayed). 3 representative cells of each class are shown. All images are rescaled the same and are of the same size scale. (b) Response class prediction of low- and high-responder cells exposed to 6.25ng/ml EGF. Logistic regression was used as classifier. The class membership of low- and high-responder was used as dependent variable. Principal components of the cellular state variables were used as independent variables. Cells were randomly assigned to the training (balanced classes) and test set. The trained model was then used to predict the class membership of the test set. Training and prediction were performed 10^4^ times and final class prediction for each cell is determined by whichever class prediction occurred the most often. (c) Predictive features for qualitative behavior (i.e. low- and high-responder) differences of pERK response at 6.25ng/ml EGF in representative cells for both response classes. Each image displaying the same marker is rescaled the same. Scale bar indicated in white. (d) The qualitative difference in pERK response behavior of individual cells is explained well only by the multivariate cellular state and not by univariate cellular features. Logistic regression models were trained with each individual feature alone and on the principal components determined by the combined features as independent variables. The class membership of low- and high-responder were used as dependent variable. Cells were randomly assigned to the training (balanced classes) and test set. Training and prediction were performed 10^4^ times. F-score of each model was calculated for high- and low-responder cells. The resulting F-score for each feature or the multivariate set is the average of the 10^4^ iterations. (e) Prediction of the continuous log_2_ pERK response at 100ng/ml in single cells. Linear regression was used to make predictions. Cells were randomly assigned to the training and test set. Training and prediction were performed 10^4^ times. Values for predicted pERK represent the mean of those iterations. Red dashed line represents the 95% observation confidence interval. Gray dots represent individual cells. (f) Side-by-side projection of measured and predicted single-cell log_2_ pERK abundance within representative populations for all concentrations of EGF. (g) Signaling response heterogeneity explained by the multivariate cellular state across all response markers and EGF concentrations. Linear regression was used to make predictions. Cells were randomly assigned to the training and test set. Training and prediction were performed 10^4^ times. Mean of the iterations was taken as final prediction and the explained variance (R^2^) calculated using those values.

Heterogeneity in signaling responses are not only of qualitative nature (e.g. multiple peaks), but display substantial quantitative variation between individual cells of up to 15-fold (**Fig. 1e; Extended Data Fig. 1g**). We thus asked to what extent the cellular state can explain the entire spectrum of responses of individual cells. Testing this on pERK at 100 ng/ml EGF, we found that its continuous response in individual cells can be well predicted (R^2^=0.84) (**Fig. 2e**). Comparing single-cell predictions with the actual measurements projected side-by-side onto the same cell population reveals the accuracy of predicting the heterogeneous response of cells exposed to the same amount of growth factor and for each concentration (**Fig. 2f**). Performing this for all 10 response markers across all 5 concentrations of EGF revealed that this generally applies, reaching coefficients of determination (R^2^) between 0.40 and 0.85 (median: 0.71) (**Fig. 2g**). We thus conclude that the heterogenous signaling response of individual mammalian cells can be largely explained by pre-existing state properties of each cell, indicating that intracellular signal processing is aware of the cellular state and context.

## Non-redundant sensing of cellular state

Signaling proteins interact in complex networks and their activation depends on each other through cascades and feedbacks^17^. It is therefore possible that the link between a heterogenous response and the cellular state could originate at one (or a few) nodes in a network, from which it is propagated throughout the rest of the network. Another possibility is that each response node, despite biochemical dependence on others, is influenced by non-redundant properties of the cellular state. In the first scenario, the activation level of most nodes encodes only shared information about the cellular state (**Fig. 3a**, dark yellow arrows) while in the latter the activation degree of each response node encodes partially non-redundant information about the cellular state (**Fig. 3a**, dark green arrows). The latter scenario might be more comprehensive as it distributes integration of information across more nodes which could each be modulated by unique aspects of the cellular state. To distinguish between these scenarios, we first looked at the regression coefficients of the cellular state models of MEK and ERK, a classical interaction pair^45^ (**Fig. 3a; Extended Data Fig. 3a**). This shows that the various aspects of the cellular state do not to the same extent predict their single-cell responses (**Fig. 3a**), supporting the latter scenario. To quantify this, we analyzed how much of the heterogeneous response in one signaling node caries unique information about the cellular state in comparison to the other signaling node, and to what extent their responses share similar information about the cellular state. In addition, we quantified the amount of information each node contains of the other node that is not linked to the cellular state (**Fig. 3a**, pink arrows). This reveals that most of the heterogeneity in pMEK and pERK is explained by the cellular state (**Fig. 3a**, bar graphs), of which a considerable fraction is unique. Thus, even in a classical interaction pair such as MEK and ERK each transmit non-redundant information about the cellular state when activated^46, 47^. We next extended this analysis to all response markers. Comparing their regression coefficients across cellular state properties showed that the heterogeneity in each response marker is explained by different aspects of the cellular state, and that collectively, they connect to a broad range of cellular state properties (**Fig. 3b**). Pairwise comparisons revealed that across a network, signaling nodes carry in part shared information, as well as unique, non-redundant information about the cellular state (**Fig. 3c; Extended Data Fig. 3b**). Signaling-specific variation between nodes that is not explained by the cellular state is usually small, and limited to directly interacting, or proximate nodes in the signaling network. For instance, while some of the heterogeneity in the response of S6 phosphorylation (pS6) is uniquely linked to variability in the co-response of pRSK, a kinase that directly phosphorylates S6^48^, as well as to the co-response of pMTOR, which achieves this via S6 kinase^49, 50^, the heterogeneity explained by the cellular state still constitutes a substantial fraction. Similar effects are seen between pAKT and pGSK3B, and the FoxO transcription factors, which are direct targets of pAKT^51, 52^ (**Fig. 3c**, highlighted; **Extended Data Fig. 3b**). This holds true even when the response of each node is compared to the combined responses of all other nodes in the signaling network (**Extended Data Fig. 3c**), indicating that each response marker is to some extent affected by different properties of the cellular state.

**Fig. 3.**
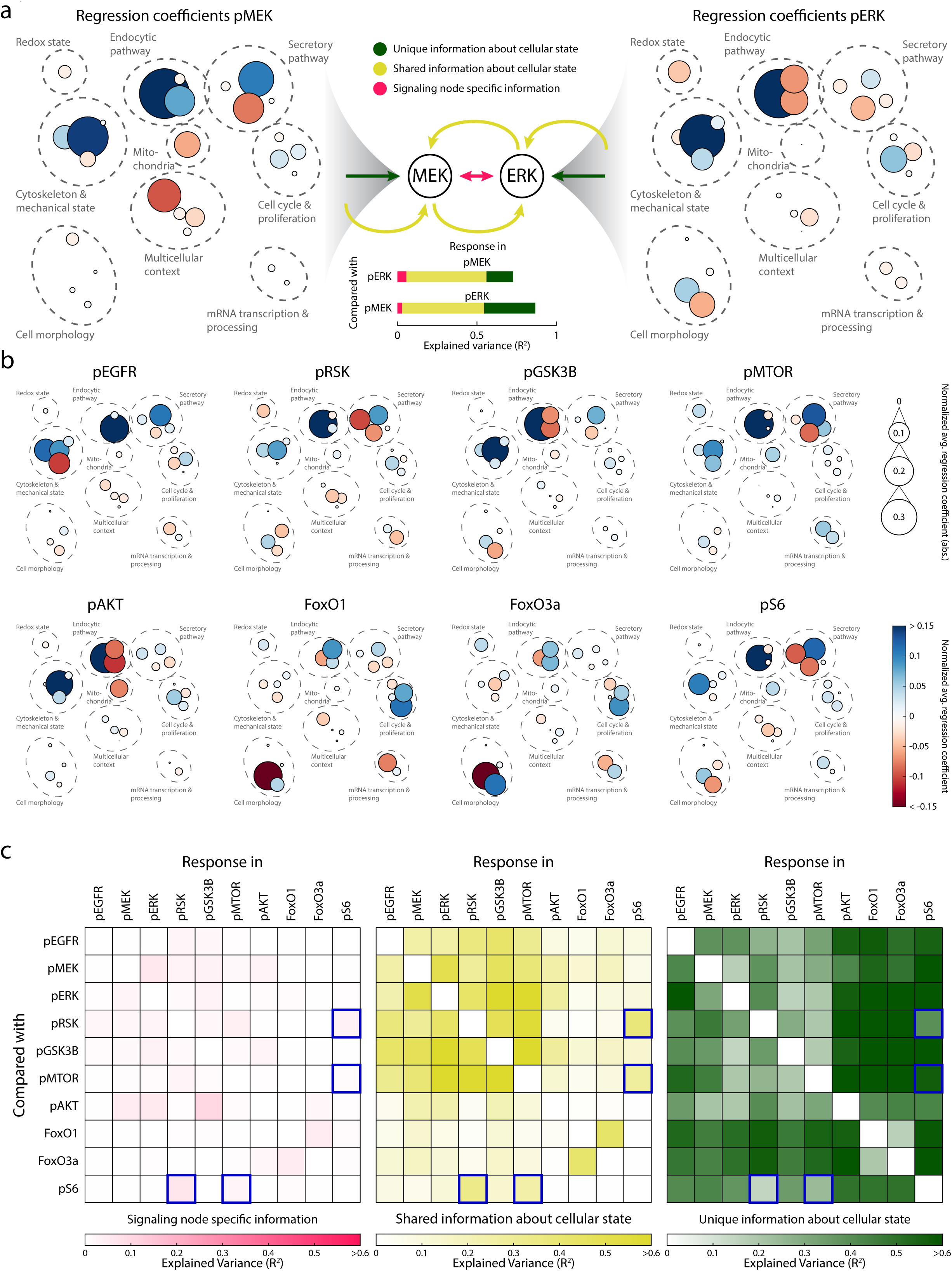
Heterogenous signaling responses carry unique information about the cellular state. (a) Manually drawn network highlighting the sum normalized, average (across all concentrations of EGF), regression coefficients (color-coded) of specific cellular features of linear models trained on the response of pMEK (left) and pERK (right). In the middle, possible scenarios of information transfer are indicated. Each node carries unique, non-redundant information about the cellular state when compared to the other (green), nodes carries shared information about the cellular state that might be propagated through their interactions (dark yellow), and each node carries additional information not related to the cellular state that is propagated through interactions (pink). Bar graphs show the respective contributions of these possible information sources and transfers in explaining the variance in signaling responses for pMEK and pERK. (b) Manually drawn networks highlighting the sum normalized, average (across all concentrations of EGF), regression coefficients (color-coded) of specific cellular features of linear models trained on the response across 8 response markers. (c) Heatmap of pairwise comparison at 100ng/ml EGF. Unique and shared fraction of variance explained of a response marker by variability in the cellular state and other response markers in displayed. Coefficient of determination (R^2^) with linear regression was calculated with three different sets of independent variables. 1: The response compared with, 2: Cellular state and 3: Comparing response combined with cellular state. Unique and specific variability explained was calculated as R^2^ difference between variable set 3 and 2 (Signaling node specific information – pink) and set 3 and 1 (Unique information about cellular state – dark green). Shared information about cellular state is calculated as variance explained by both variation in the response compared and cellular state variation (3 - (3-1) - (3-2)) (dark yellow). Pairwise comparison of the nodes pS6 with pRSK, and pMTOR are highlighted with blue boxes related to the main text.

To further explore the cellular state properties that have the strongest unique impact on each response node, we started with the response markers pERK and pMTOR, which are well-characterized kinases that have been linked to different properties of the cellular state^35, 36, 39, 40^. Plotting their single-cell abundances after 5 min of exposure to 100ng/ml EGF reveals strong correlation (Pearson’s r = 0.82). However, overlaying these correlation plots with the abundances of early endosomes (EEA1) and mitochondria (HSP60) reveals that the activation of ERK and MTOR exhibit distinct dependencies on these organelles (**Fig. 4a**). The variation in pERK not explained by pMTOR is associated more strongly with variability in EEA1 than the variation in pMTOR that is not explained by pERK (respective partial correlations: 0.51 and 0.31). In contrast, only the variation in pMTOR not explained by pERK is associated positively with variability in HSP60 (respective partial correlations: -0.14 and 0.49). This shows that pERK senses more strongly the abundance of early endosomes in individual cells than pMTOR, while pMTOR senses more strongly the abundance of mitochondria, in agreement with well-known respective connections between these organelles and signaling nodes^35, 36, 39, 40^.

**Fig. 4.**
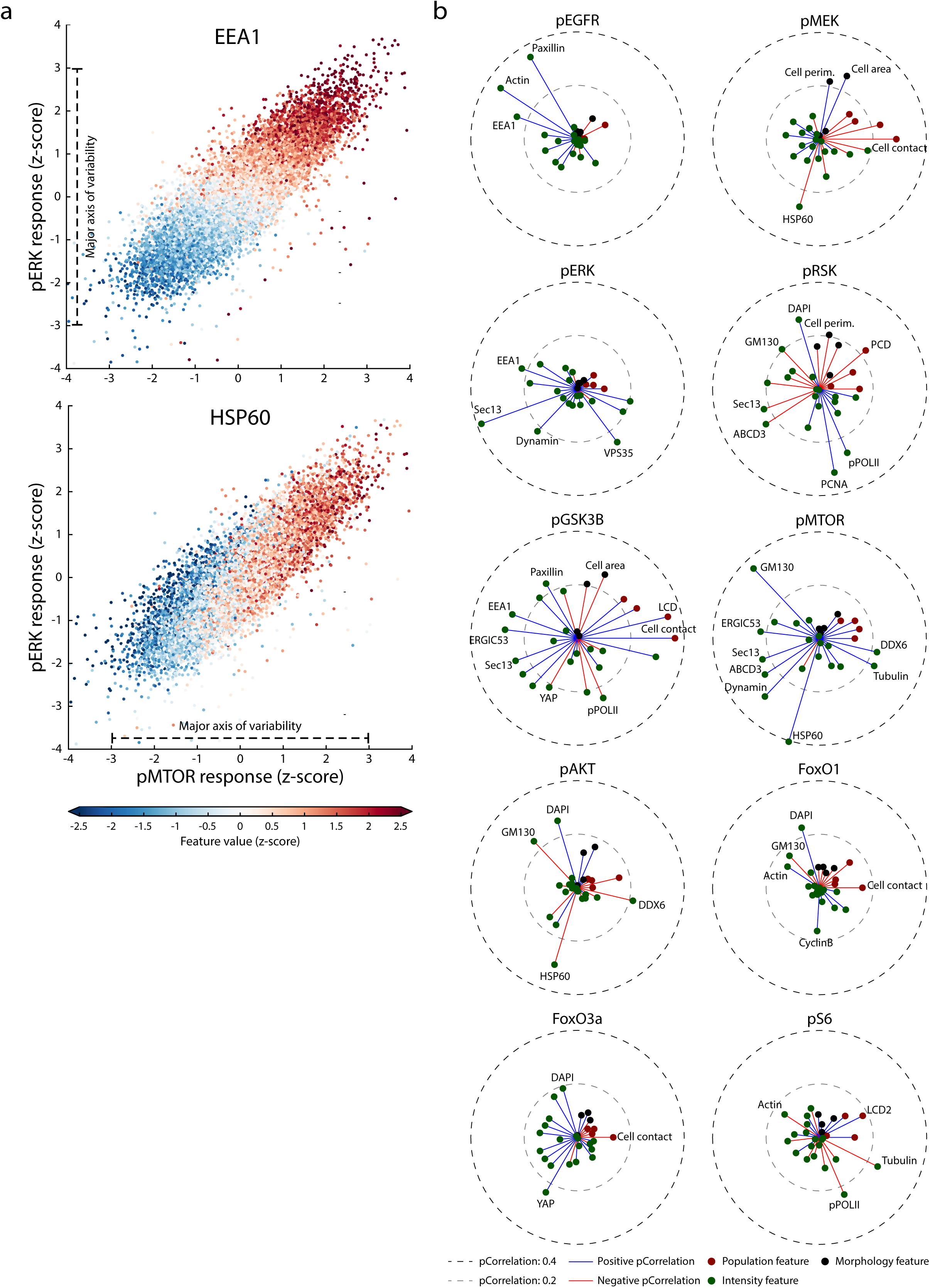
Non-redundant sensing of cellular state. (a) Scatter plot of z-scored log_2_ pMTOR mean abundance against z-scored log_2_ pERK mean abundance in single cells at 100ng/ml of EGF. In the top panel each scatter point is colored according to its z-scored mean abundance of EEA1 of each respective cell. In the bottom panel scatter point is colored according to its z-scored mean abundance of HSP60 of each respective cell. (b) Each response marker displays a unique association pattern with specific properties of the cellular state. For each response marker, partial correlations (partial to all other response markers) were calculated to a subset of cellular state features. Partial correlations for each response marker are displayed in a star plot. Manually selected features are highlighted and labeled. Partial correlations to specific features presented are from 100ng/ml EGF. Partial correlation is abbreviated to “pCorrelation”.

We extended this analysis to the whole signaling network by calculating partial correlations of each response marker (partial to all other responses) with cellular state features (**Extended Data Table 4; Fig. 4b**). This revealed response markers exhibiting non-redundant sensing of a smaller set of cellular state properties, such as pERK, pMEK and pAKT, and response markers that sense a broader set of properties, such as pRSK, pMTOR and pGSK3B. Interestingly, kinases which are directly linked to each other for activation, such as MEK, ERK and RSK, can display markedly different association patterns. For instance, the unique association pattern of pERK is predominantly linked to state markers of endocytic membrane trafficking (EEA1, VPS35, Sec13 and Dynamin), while the abundance of pMEK, its direct upstream kinase, shows unique positive association with cell area and negative association with the extent of cell-cell contact and mitochondrial abundance (HSP60). The third kinase in the cascade, RSK, displays yet another pattern of unique associations. Its activation is negatively associated with cellular crowding, cell area and cell perimeter while exhibiting positive association with features linked to cell cycle progression (DNA content and PCNA abundance) and transcriptional activity (pPOLII). For the activation of MTOR we find unique positive association with state markers of secretory membrane trafficking (ERGIC53 and GM130), mitochondrial abundance (HSP60) and the cytoskeleton (Tubulin). Overall, these association patterns are consistent between similar concentrations of EGF (**Extended Data Fig. 4a**). We conclude that signaling response markers display quantifiable unique and thus non-redundant association with specific properties of the cellular state in their acute response to EGF.

## Cellular state-dependent sensitivities to EGF

We then asked whether the cellular state also affects sensitivities of individual cells to different concentrations of EGF, by inferring dose-response relationships from groups of cells that are in the same cellular state but exposed to different amounts of growth factor obtained by fuzzy clustering and comparing them between different cellular states (**Fig. 5a; Extended Data Fig. 5a-d**). This showed that signaling responses display widely varying sensitivities and dynamic ranges to EGF depending on the cellular state. To exemplify this, we depicted the level of pERK of groups of cells in similar cellular states relative to the mean of the whole population (**Fig. 5b**). For pERK, a lower then average response at low concentration of EGF is, amongst others, associated with high local cell density (cellular state 2 and 12), whereas a lower than average response at high concentration of EGF is, amongst others, associated with a low abundance of early endosomes (EEA1) (cellular state 12 and 15) (**Fig. 5a**, dashed boxes). Extending this analysis to all signaling nodes by calculating cellular state-dependent half-maximal effective concentrations (EC50) of EGF revealed similarly strong effects of the cellular state on the EC50 of each signaling response (**Fig. 5c**). This indicates that activation of each signaling node exhibits state-aware sensitivity to the spectrum of EGF concentrations cells are exposed to.

**Fig. 5.**
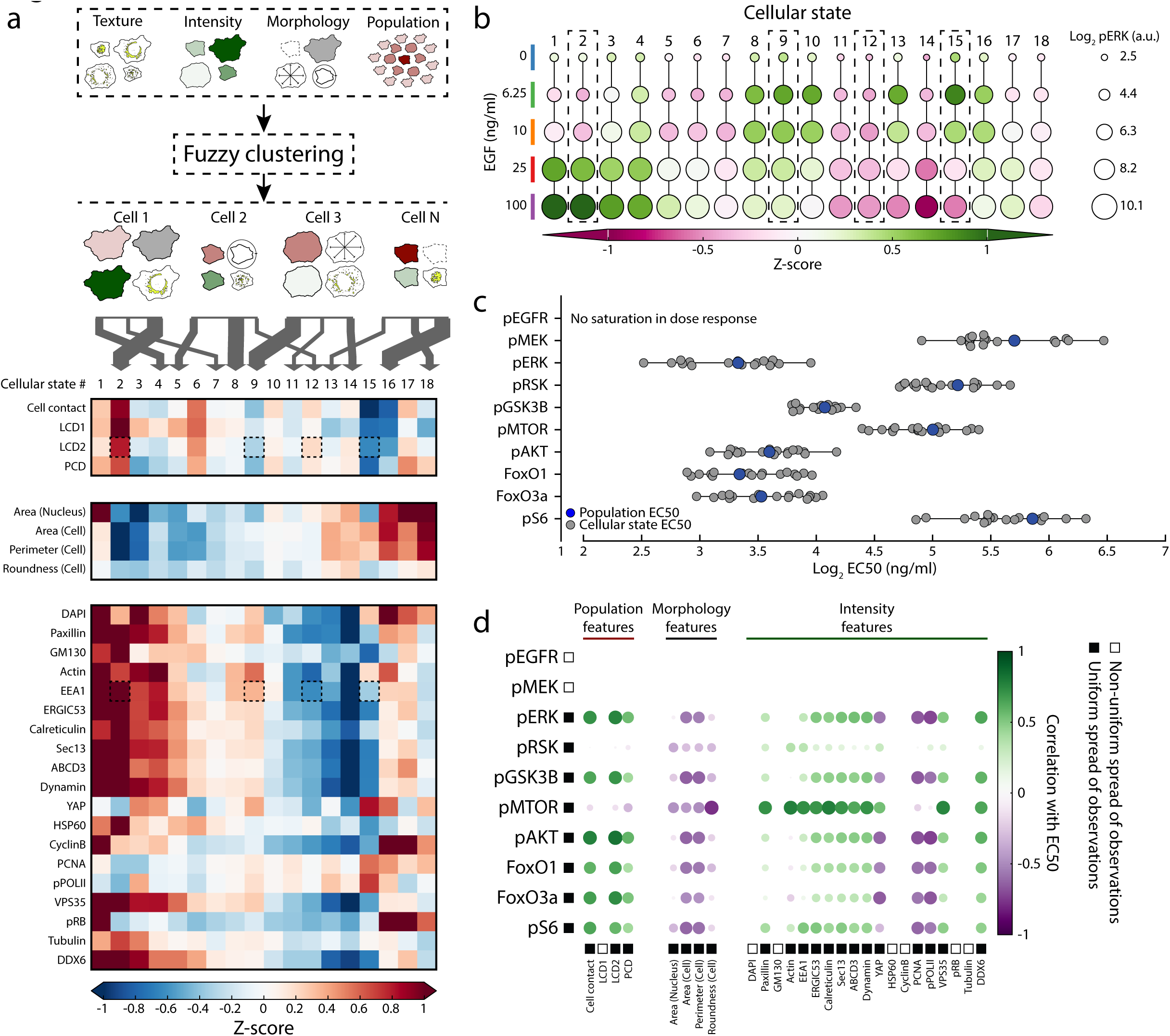
Cellular state-dependent sensitivities to EGF. (a) Schematic representation of and results from c-means (fuzzy) clustering of cells in the multivariate cellular state space. Fuzzy clustering assigns each cell a membership degree to each cluster centroid biased by its distance from it. Membership degree of each cells sums to 1. Signaling relevant principal components (PC) were identified by stepwise elimination of PCs in linear regression on signaling response markers across all concentrations of EGF. PCs found as significant predictor in at least 90% of all models were used for clustering. Fuzzy clustering with Manhattan distance was performed 10^4^ and the result with the lowest objective function was retained. Heatmap depicts the weighted mean (weighted by membership degree of each) feature value (z-scored) of a readily interpretable subset of cellular state features. Dashed boxes highlight selected features in specific cellular states. (b) Bubble plot of mean log_2_ pERK abundance in each fuzzy cluster across concentrations of EGF. Size of circles represent the raw value of mean log_2_ pERK abundance while colors depict the mean z-score (z-scored in each concentration of EGF) of the mean log_2_ pERK abundance in each cluster. Dashed boxes highlight the pERK response in specific cellular states. (c) Half maximal effective concentrations (EC50) of EGF for each response marker and cellular state. EC50 was calculated using a 4-parameter logistic fit to the normalized (0-1) response across concentrations of EGF in each cluster. Gray dots represent values obtained for individual cellular states. Blue dots represent values obtained from the whole, non-stratified, population of cells. (d) Bubble plot of correlation analysis for EC50 values and specific properties of the cellular state using the stratified samples. Features and EC50 values which do not exhibit a uniform spread across the range of values were excluded for correlation analysis to obtain reliable estimates.

Measuring the correlation of individual state features with the EC50 values, could reliably identify features that render cells more sensitive (negative correlation) or less sensitive (positive correlation) to EGF for 8 of the 10 signaling nodes (**Fig. 5d**; **Extended Data Fig. 5e**). Six of them (pERK, pAKT, pGSK3B, FoxO1, FoxO3a, and pS6) revealed a similar pattern of associations, in which their sensitivities to EGF decreased with increasing local cell density and cell contacts while they increased with increasing cell area, cell perimeter and higher transcriptional activity (pPOLII). A different type of pattern was displayed by pMTOR and pRSK, whose sensitivities to EGF did not exhibit a dependency on local cell density or the amount of pPOLII, but increased with increasing cell and nuclear area, suggesting that the latter two are specifically adapted to cell size independent of population context or transcriptional activity. Moreover, the sensitivity of pMTOR to EGF decreases strongly with increased abundance of intracellular organelles suggesting an adaptation of EGF-induced cellular growth to cellular content. Taken together, this shows that signaling nodes exhibit sensitivities adapted to the cellular state across a spectrum of EGF concentrations.

## Multimodal signaling enables accurate information processing

Having established that signaling nodes show profound non-redundant sensing of different aspects of the cellular state and that this influences their sensitivity to EGF, points to an intracellular equivalent of multimodal perception. Here, multiple signaling nodes (modes) of a network would enable an individual cell to sense its position within the complex landscape of cellular states and accurately place EGF concentration within this context. To test the former, we analyzed the similarity in cellular states amongst cells that exhibit similar signaling responses. This showed that individual responses are substantially less informative about similarity in cellular state than the multimodal response. (**Fig. 6a**). To test the latter, we predicted the concentration of EGF an individual cell is exposed to from its acute signaling response. Predictions based on only the abundance of pERK, or any of the other individual response markers, performed, as expected, not much better than a random guess (**Fig. 6b; Extended Data Fig. 6a**). However, when the pERK signaling response of individual cells was placed within the context of the cellular state by incorporating cellular state features, predictions improved substantially (**Fig. 6c**). Importantly, features of the cellular state alone did not have any predictive power since they do not change during a 5 min exposure to EGF (**Extended Data Fig. 1e; Extended Data Fig 6b**). Nevertheless, some ambiguity still remained for higher concentrations of EGF. Since activation of ERK is particularly sensitive to changes in low concentrations of EGF, we next included the other signaling nodes, as these may provide additional information on changes in higher concentrations. This showed that the multimodal signaling response based on all response markers eliminated most uncertainty and achieved a near-perfect prediction (median: 0.90) of the correct concentration for each individual cell (**Fig. 6d**). This was only achieved when the multimodal signaling response was considered within the context of each cell’s cellular state (**Extended Data Fig. 6c**). Thus, individual cells have the capacity to accurately distinguish distinct concentrations of EGF by means of non-redundant multimodal cellular state-aware signaling.

**Fig. 6.**
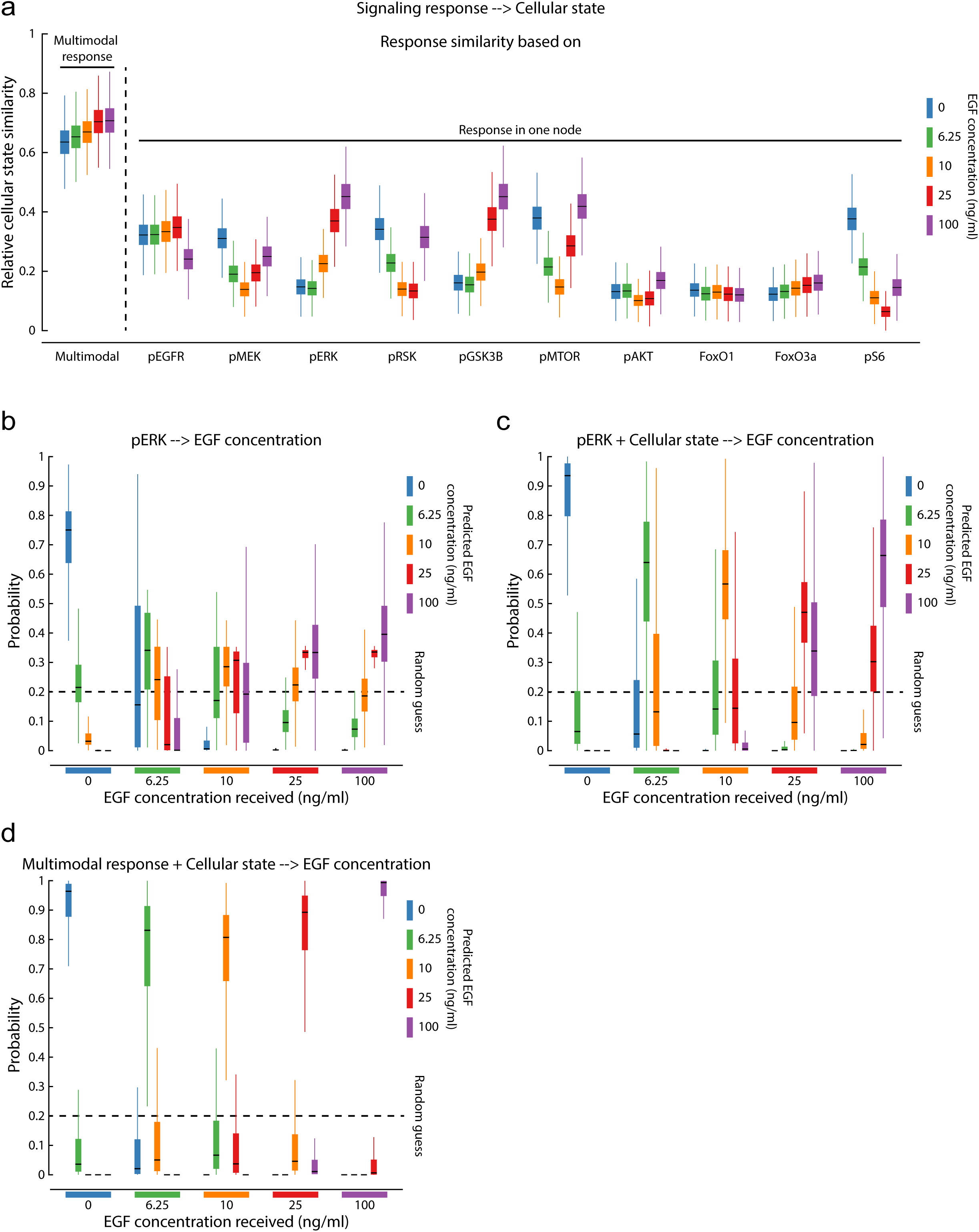
Multimodal signaling enables accurate information processing. (a) Neighbor (of response) similarity (of cellular state) comparison of the multimodal response and each individual response marker alone. For each cell the closest 50 neighbors in multimodal response or single response space (each marker individually) were calculated using Manhattan distance. Then 20 cells were randomly selected 10^5^ times and the mean cosine similarity in cellular state space (readily interpretable subset) for each cell’s closest neighbors calculated in both multimodal response and single response space. Values are then normalized by the highest obtained similarity value. Boxplots display the distribution of similarities in cellular state space obtained from bootstrapped sampling for each concentration EGF. Boxplots for each single response marker and multimodal response are displayed together. (b) Boxplots of predicted EGF concentration received. A multiclass logistic classifier (Concentration as classes) with balanced classes was trained with single-cell log_2_ pERK mean abundances as independent variable. Model was then used to predict the probability per class on a separate set of single cells and displayed separated by true class. Model prediction was performed with random subset of training cells and test cells 150 times or till 10 predictions per single cell was obtained. Displayed probability for each cell is the mean of all single predictions. Dashed line represents the random guess probability. (c) Boxplots of predicted EGF concentration received. A multiclass logistic classifier (Concentration as classes) with balanced classes was trained with single-cell log2 pERK mean abundances and the scale-crossing cellular state as independent variables. Model was then used to predict the probability per class on a separate set of single cells and displayed separated by true class. Model prediction was performed with random subset of training cells and test cells 150 times or till 10 predictions per single cell was obtained. Displayed probability for each cell is the mean of all single predictions. Dashed line represents the random guess probability. (d) Boxplots of predicted EGF concentration received. A multiclass logistic classifier (concentration as classes) with balanced classes was trained with all response markers combined (multimodal signaling) and the scale-crossing cellular state as independent variables. Model was then used to predict the probability per class on a separate set of single cells and displayed separated by true class. Model prediction was performed with random subset of training cells and test cells 150 times or till 10 predictions per single cell was obtained. Displayed probability for each cell is the mean of all single predictions. Dashed line represents the random guess probability.

## Discussion

We here quantified, *in situ* and across multiple spatial scales, acute changes in the activated state of 7 kinases, 2 transcription factors, and 1 ribosomal subunit upon exposure to EGF, along with 18 non-changing markers of the cytoskeleton, membrane trafficking, cytoplasmic RNA processing, and the mechanical, proliferative, metabolic, and transcriptional state simultaneously in individual cells. The resulting dataset reveals a strong link between the cellular state and the responses of individual cells to an incoming signal. This effect arises from multiple spatial scales, and collectively explains a large fraction of heterogeneity observed in single-cell responses to growth factor exposure. It shows that signaling nodes in a network are not equally linked to the same cellular properties, and that the variability in output response to an equal input displayed by individual nodes encodes partially non-redundant information. Collectively, the non-redundant information contained in the multimodal response is a more accurate representation of a cell’s cellular state than each individual response alone. In addition, individual nodes exhibit cellular state-dependent sensitivity ranges across different input concentrations of growth factor. This suggests that the multimodal, cellular state-dependent nature of signal transduction in mammalian organisms has evolved to endow individual cells with a high capacity for information processing to accurately interpret a broad range of input information within the complex landscape of cellular states and take the appropriate decision. Interestingly, the properties of intracellular signal processing revealed here are similar to multimodal perception performed by sensory systems across the kingdom of multicellular life^31, 53–56^. Both display sensing through multiple modalities (multiple signaling nodes versus multiple sensory neurons) with partial non-redundancy, which allows the integration of multiple sources of information into a more complete picture of reality. Both also display cross-modal effects where activation of one mode can influence the perception in another^57–63^ (e.g. the ventriloquist effect versus crosstalk in signaling networks). In addition, the partial redundancy of nodes makes cellular state sensing robust to noise or errors in the signal of individual nodes, similar to the role of partial redundancy in information content between auditory and visual perception. This adds to the growing notion that biological systems operate according to similar rules despite occurring at vastly different scales.

While our study focused on the acute response 5 min after addition of growth factor to avoid EGF-induced changes in the cellular state, our approach could in the future be extended to time courses by updating the cellular state properties as cells progress in time. This may cast a new light on well-known characteristics of intracellular signaling dynamics. It may reveal whether differences between sustained versus transient or repetitive (e.g. pulsatile and oscillatory) responses in individual cells can also be explained by the cellular state, as similar mechanisms that make the activation of signaling nodes dependent on the intracellular context likely also apply to their inactivation. This can lay the basis for identifying the molecular mechanisms underlying multimodal perception in individual cells, in which certain cellular state properties sensed by one signaling node may, via connections between nodes, lead to sustained activation, or rapid deactivation of another signaling node that senses other state properties, creating a percept that is unique to the specific combination of cellular state properties of that cell. Moreover, it may reveal a new type of feedback that crosses scales, in which acute responses lead to changes in the cellular state, which in turn influence the signaling response. Different dynamic properties, which can translate to different downstream responses^15, 16, 19, 64–66^, may thus reflect the mechanisms by which multivariate information about the cellular state is transmitted to cellular decision making. Despite a large body of work on specific mechanisms by which cellular structures influence signaling, the cellular state remains a largely unexplored source of information in our quantitative understanding of life. We expect that by integrating the cellular state into the analysis of single-cell decision making in development, as well as of single-cell drug responses in cancer^67^ and other diseases as shown here, predictability will increase to levels hitherto inconceivable.

## Acknowledgements

We thank all members of the Pelkmans lab for discussions. We further thank Robin Klemm, Prisca Liberali and Wilfried Kramer for critical reading of the manuscript. L.P. is supported by the Swiss National Science Foundation and the University of Zurich.

---

**Extended Data Fig. 1.**
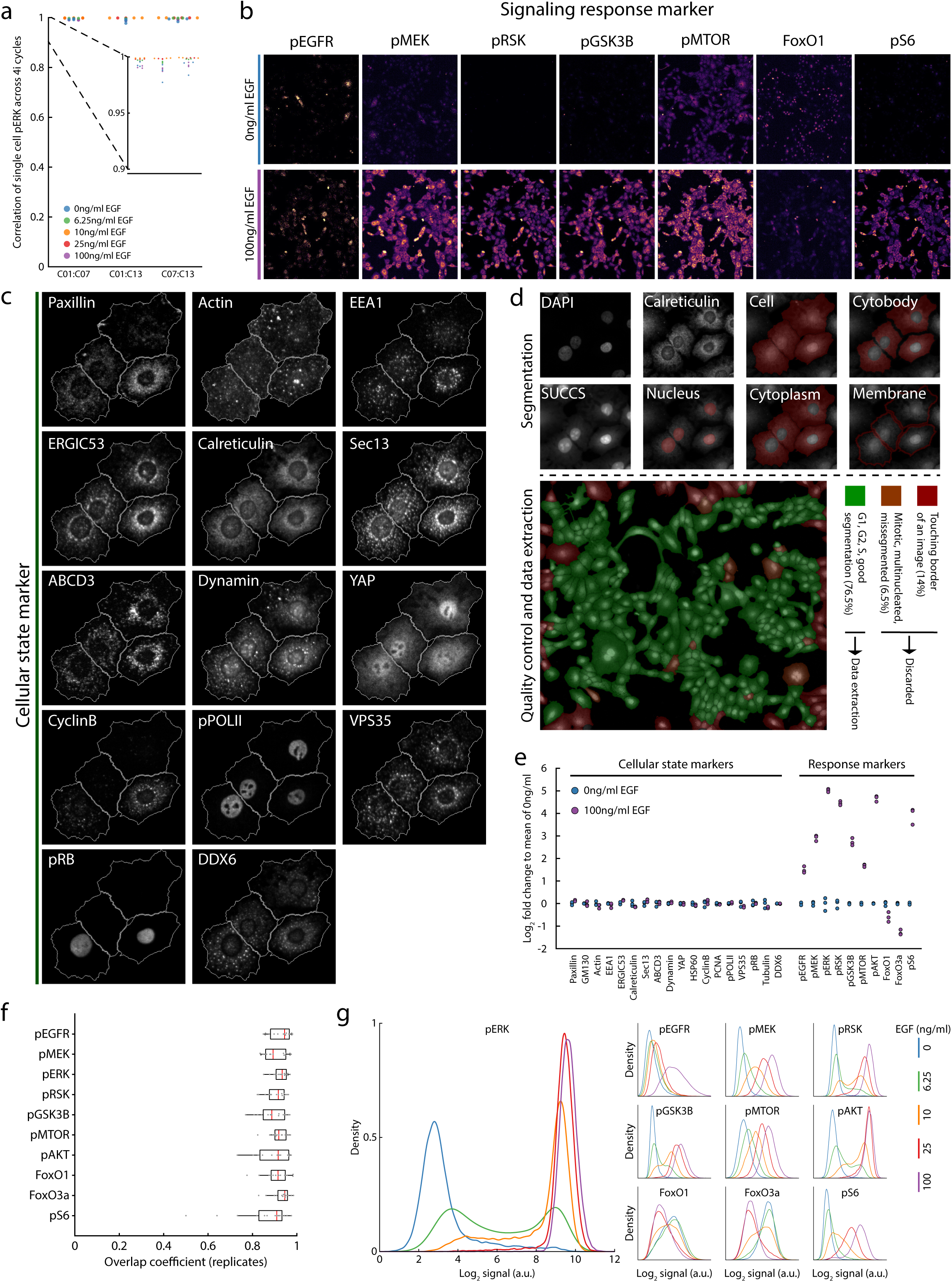
4i allows reproducible quantification of acute changes in signaling response and the non-changing cellular state simultaneously in thousands of individual cells. (a) 4i achieves highly reproducible quantification of activated signaling proteins across large numbers of single cells. Repeated staining against phosphorylated ERK in different cycles (01, 07 and 13) across 5 concentrations of EGF in 3 replicates each. Pearson correlation coefficient is calculated for single-cell, cytoplasmic abundances of pERK across all cycle combinations and conditions. Dashed lines highlight a zoom-in of the indicated axis. (b) Visualization of response markers obtained by 4i in representative populations exposed to either 0ng/ml or 100ng/ml EGF for 5 min. Same population displayed for each response marker and condition. (c) Visualization of state markers obtained by 4i in representative cells. Same cells displayed for each state marker. (d) Schematic representation of the object segmentation and quality control pipeline. Original images and segmentation objects are shown for 4 representative cells (above dashed line). Segmentation is performed on DAPI (cycle 01), SUCCS (cycle 20) and Calreticulin (cycle 12) images. 5 distinct regions are segmented and highlighted in red on the composite of DAPI, SUCCS and Calreticulin. Quality control is then performed and depicted for one representative imaging site (below dashed line). Miss-segmented, multinucleated and mitotic cells (highlighted in orange) and cell segmentations touching the boundaries of an image acquisition site (highlighted in red) are excluded. The remainder of cells is retained (highlighted in green) and features are extracted. (e) Response marker display a robust change in abundance upon EGF stimulation whereas state markers do not. Log_2_ fold change of each replicate to the mean of all unstimulated replicates was calculated. Dots represent individual replicates. (f) Response marker distributions are reproducible across independent replicates. Overlap coefficient (shared area under two probability density function) for each combination of replicates and for each concentration was calculated using Monte-Carlo integration and is displayed. Empirical probability density functions where used and are calculated using kernel density estimation with a gaussian kernel. (g) Heterogenous single-cell response give rise to overlapping distributions across EGF concentrations. Value density inference was performed using kernel density estimation (gaussian kernel, bandwidth chosen according to Silverman’s rule of thumb) on log_2_ transformed values for each response marker and each concentration of EGF. Each subpanel depicts a different response marker and the associated density estimates for each concentration of EGF.

**Extended Data Fig. 2.**
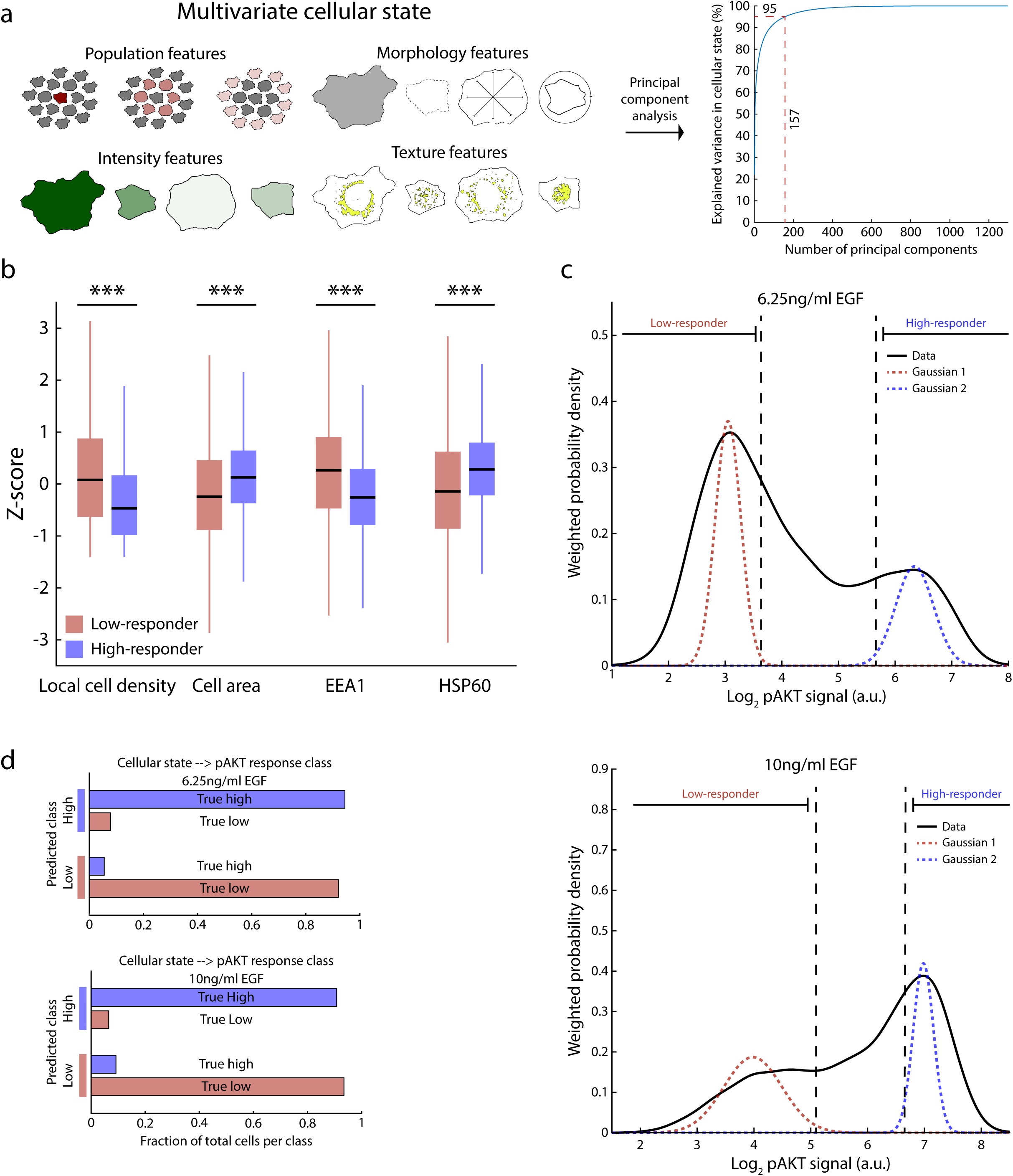
Cellular state determines the heterogeneous signaling response. (a) Transformation of variables describing the multivariate cellular state into uncorrelated principal components. PCA was performed on the complete data-set and the cumulative variance explained is depicted as a function of number of principal components included. 95% cumulative variance explained was chosen as cutoff resulting in 157 principal components describing the cellular state. (b) Individual features that allow the prediction of pERK behavior arise at multiple spatial scales. Boxplot of z-scored feature values from 4 features from distinct spatial scales of low- and high-responder cells are displayed. Two-sample Kolmogorov-Smirnoff test was performed against the null hypothesis that both data are from the same continuous distribution. For all tested features the null hypothesis was rejected at significance level of 0.001 (***). (c) Phosphorylation of AKT in cell populations exposed to 6.25ng/ml (top panel) or 10ng/ml (bottom panel) of EGF exhibited two distinct peaks. Kernel density estimates with a gaussian kernel of the data are displayed. Cells were separated into low- and high-responder cells based on a gaussian mixture model (2 Gaussian components displayed). (d) Response class prediction of low- and high-responder cells exposed to 6.25ng/ml EGF (top panel) or 10ng/ml EGF (bottom panel). Logistic regression was used as classifier. The class membership of low- and high-responder was used as dependent variable. Principal components of the cellular state variables were used as independent variables. Cells were randomly assigned to the training (balanced classes) and test set. The trained model was then used to predict the class membership of the test set. Training and prediction were performed 10^4^ times and final class prediction for each cell is determined by whichever class prediction occurred the most often.

**Extended Data Fig. 3.**
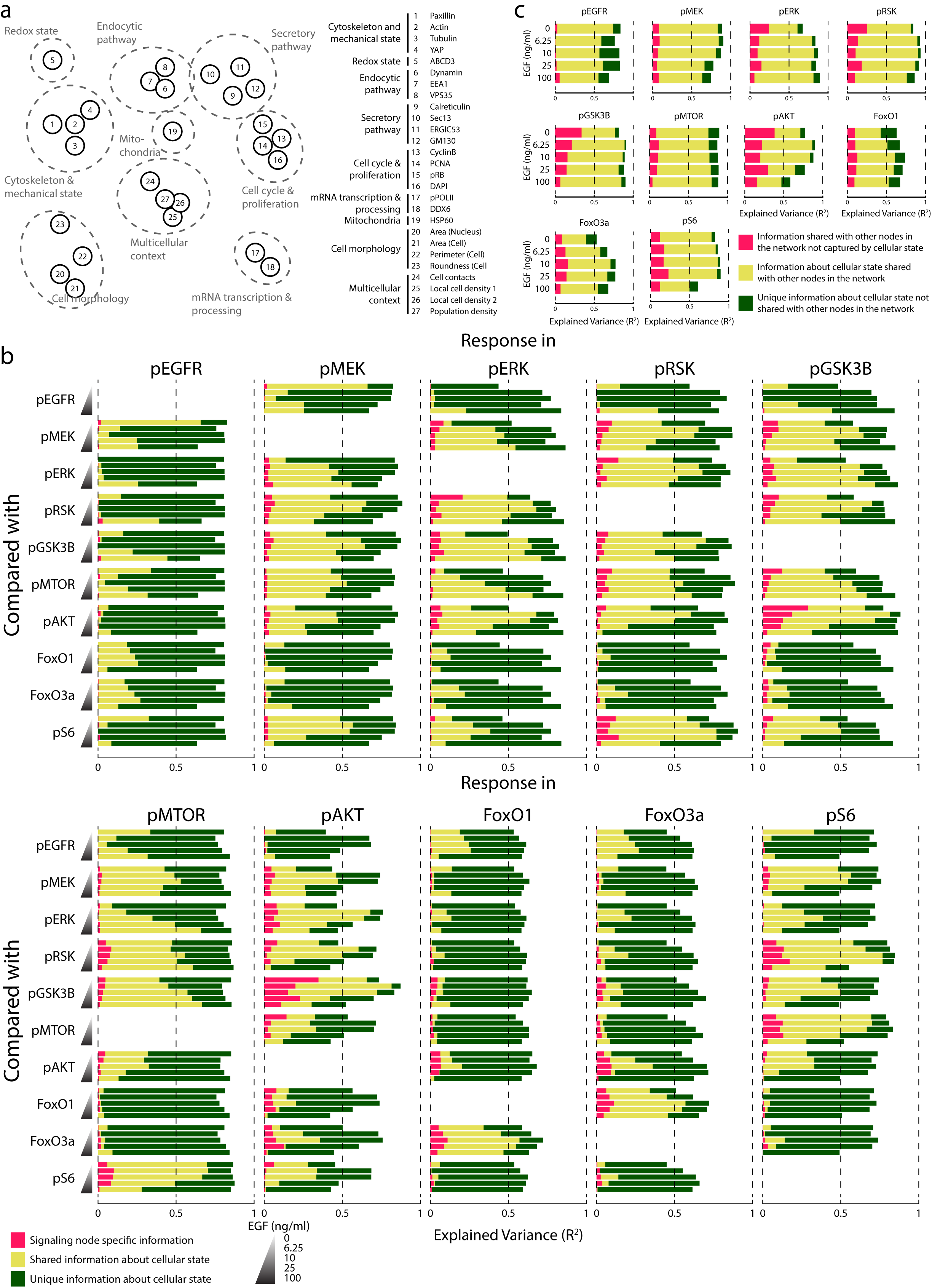
Heterogenous signaling responses carry unique information about the cellular state. (a) Schematic representation of a prior-knowledge based ordering of cellular state features into a network. Nodes represent individual features and are highlighted by gray circles corresponding to their functions. Table lists the names and assignment for every feature highlighted. (b) Stacked bars of pairwise comparison across all concentrations of EGF. Unique and shared fraction of variance explained of a response marker by variability in the cellular state and other response markers is displayed. Coefficient of determination (R^2^) with linear regression was calculated with three different sets of independent variables. 1: The response compared with, 2: All other responses 3: All other responses and cellular state. Unique variability explained was calculated as R^2^ difference between variable set 3 and 2 (signaling node specific information – pink) and set 3 and 1 (Unique information about cellular state – dark green). Shared information about cellular state is calculated as variance explained by both variation in the compared response and cellular state variation (3 - (3-1) - (3-2)) (dark yellow). (c) Stacked bars of comparison against all other responses across all concentrations of EGF. Unique and shared fraction of variance explained of a response marker by variability in the cellular state and other response markers is displayed. Coefficient of determination (R^2^) with linear regression was calculated with three different sets of independent variables. 1: All other responses 2: Cellular state and 3: All other responses and cellular state. Unique variability explained was calculated as R^2^ difference between variable set 3 and 2 (signaling node specific information – pink) and set 3 and 1 (Unique information about cellular state – dark green). Shared information about cellular state is calculated as variance explained by both variation in the compared responses and cellular state variation (3 - (3-1) - (3-2)) (dark yellow).

**Extended Data Fig. 4.**
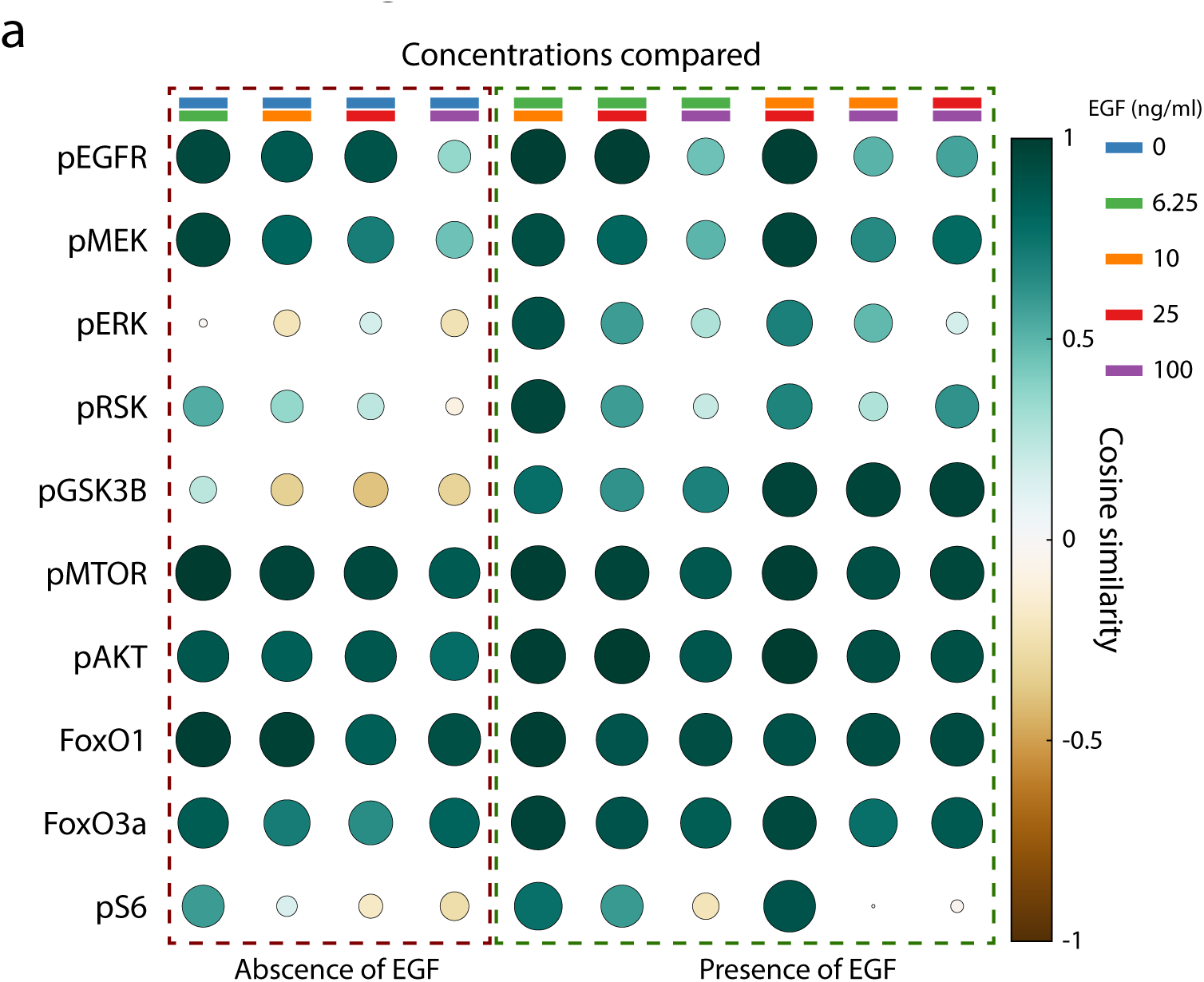
Unique association patterns of signaling responses with cellular features are consistent across concentrations of EGF. (**a**) Similarity of unique association patterns with cellular state features for each response marker across concentrations of EGF in a pairwise comparison. For each response marker, partial correlations (partial to all other response markers) were calculated to a readily interpretable subset of cellular features in each concentration of EGF. The obtained partial correlation profiles of each response marker were then compared across concentrations of EGF by calculating the cosine similarity of these resulting association patterns.

**Extended Data Fig. 5.**
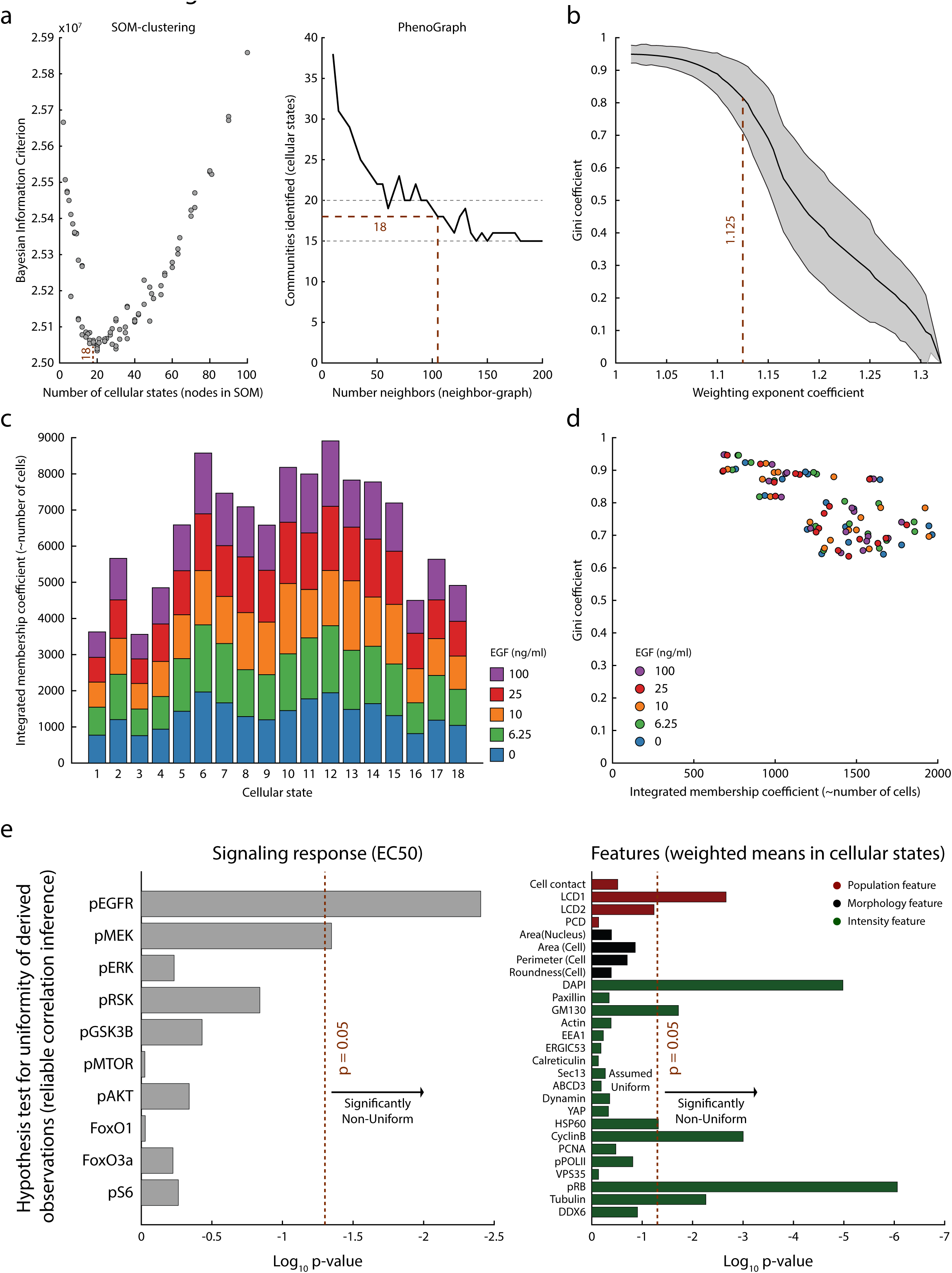
Reliable estimates of EGF sensitivity and its association with cellular state features can be obtained by means of fuzzy clustering. (a) Heuristic for choosing good number of centroids for c-means (fuzzy) clustering. Left plot depicts estimation using Self-Organized Maps (SOM). The principal components describing the multivariate cellular state of each cell were subjected to clustering by SOMs across different sized initial grids of neurons (i.e. number of clusters). The Bayesian Information Criteria (BIC) was then calculated on the results for each clustering. 18 number of centroids was identified close to the minimum in BIC. Right plot depicts centroid estimation using PhenoGraph clustering. PhenoGraph analysis was performed across different numbers of closest neighbors used to build the initial Neighbor-Graph. Communities (i.e. number of clusters) identified displayed a plateau at 20 and 15 identified communities (gray dotted line) with 18 representing the middle of slope between both plateaus. (b) Heuristic for choosing a good value of the weighting exponent coefficient (WEC) used in c-means clustering. The c-means clustering output is a vector of membership degrees for each cells and each cluster. The “fuzziness” can be modulated by the WEC. For the analysis a good consensus between biased yet non-discrete membership degrees were preferred. Gini-coefficient of membership degree values was calculated for each cluster. Mean value is depicted as black line and standard deviation is depicted as grey patch as function of the WEC. (c) Cluster membership degrees for each fuzzy cluster are not biased by concentration EGF. Integrated membership degrees of cells for each cluster separated by concentration of EGF are displayed as stacked bars. (d) A good consensus between fuzziness and biased membership degrees is obtained for each cluster across all concentrations of EGF. Gini coefficient was calculated for each cluster and each concentration of EGF and is depicted as scatter plot against the integrated membership degree. (e) Derived EC50s and feature values in each cluster are largely uniformly spread. Hypothesis testing against the null hypothesis that values are samples from a uniform distribution was performed by one-sample Kolmogorov-Smirnov test. Null hypothesis was rejected at p = 0.05 (depicted as red dashed line).

**Extended Data Fig. 6.**
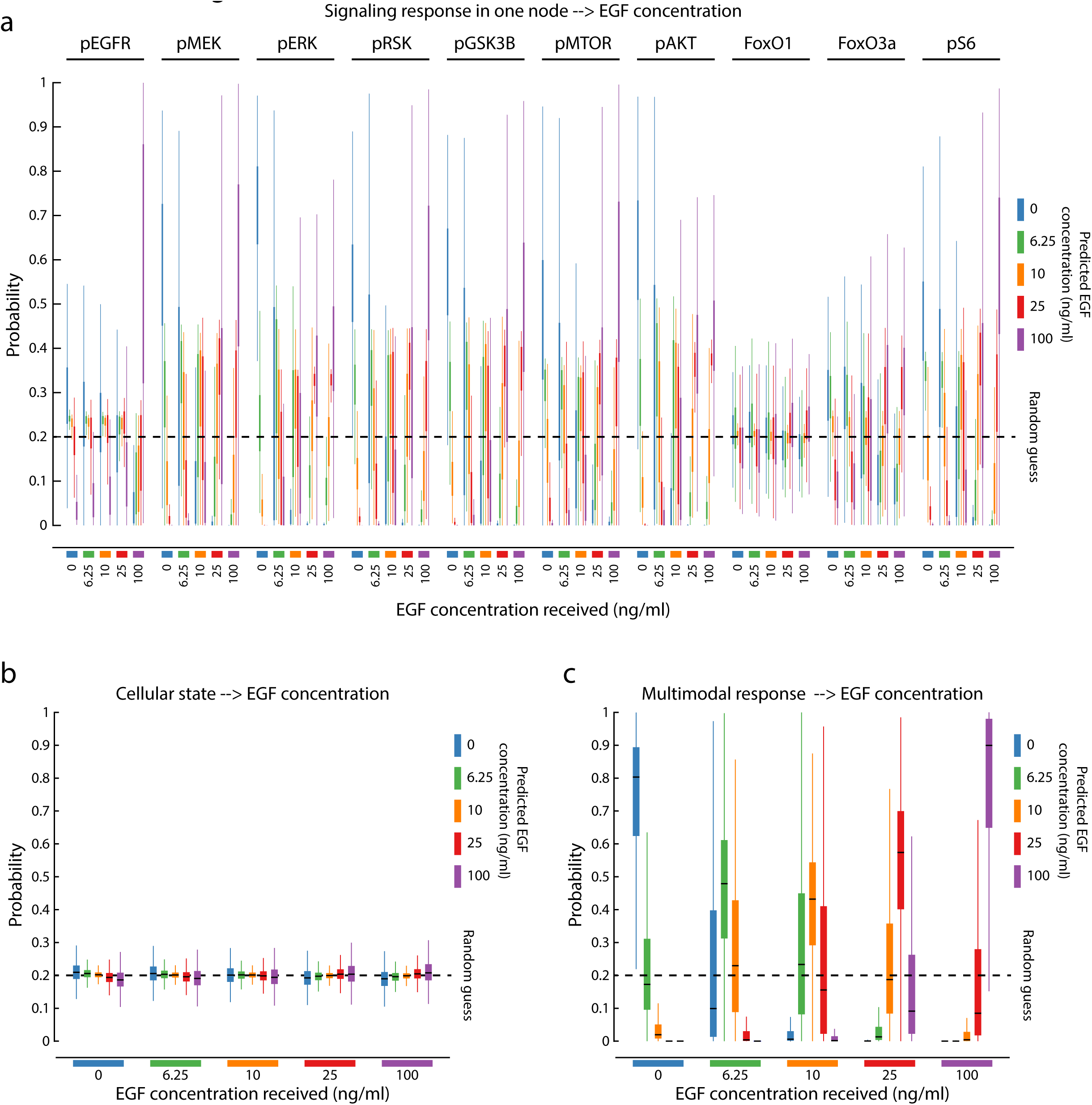
Multimodal signaling enables accurate information processing. (a) Boxplots of predicted EGF concentration received. A multiclass logistic classifier (Concentration as classes) with balanced classes was trained with each response marker individually as independent variable. Model was then used to predict the probability per class on a separate set of single cells and displayed separated by true class. Model prediction was performed with random subset of training cells and test cells 150 times or till 10 predictions per single cell was obtained. Displayed probability for each cell is the mean of all single predictions. Each response marker is displayed individually. Dashed line represents the random guess probability. (b) Boxplots of predicted EGF concentration received. A multiclass logistic classifier (Concentration as classes) with balanced classes was trained with the cellular state features as independent variables. Model was then used to predict the probability per class on a separate set of single cells and displayed separated by true class. Model prediction was performed with random subset of training cells and test cells 150 times or till 10 predictions per single cell was obtained. Displayed probability for each cell is the mean of all single predictions. Dashed line represents the random guess probability. (c) Boxplots of predicted EGF concentration received. A multiclass logistic classifier (concentration as classes) with balanced classes was trained with all response markers combined (multimodal signaling) as independent variables. Model was then used to predict the probability per class on a separate set of single cells and displayed separated by true class. Model prediction was performed with random subset of training cells and test cells 150 times or till 10 predictions per single cell was obtained. Displayed probability for each cell is the mean of all single predictions. Dashed line represents the random guess probability.

## Materials

### Cell line

Single-cell clone derived from 184A1 (human breast epithelial cell line, ATCC® CRL-8798). Cells were tested for absence of mycoplasma before use.

### Growth factor stimulation

Epidermal Growth Factor (EGF) (Millipore).

### Complete growth medium (CGM)

CGM is composed of 5% Horse Serum (HS), 20ng/ml EGF, 10µg/ml Insulin, 0.5µg/ml Hydrocortisone, 10ng/ml Cholera Toxin in DMEM/F12. HS (Thermo Fisher Scientific), EGF (Millipore), Insulin (Sigma-Aldrich), Hydrocortisone (Merck), Cholera Toxin (Sigma-Aldrich), DMEM/F12 (Thermo Fisher Scientific).

### Assay medium (AM)

AM is composed of 0.5µg/ml Hydrocortisone, 10ng/ml Cholera Toxin in DMEM/F12. Hydrocortisone (Merck), Cholera Toxin (Sigma-Aldrich), DMEM/F12 (Thermo Fisher Scientific).

### 4i blocking solution (sBS)

sBS is composed of 150mM Maleimide and 2% Bovine Serum Albumin (BSA) in phosphate buffered saline (PBS). Maleimide is added to the aqueous solution 15 minutes before the blocking step in the 4i protocol. Maleimide (Sigma-Aldrich), BSA (Sigma-Aldrich).

### Conventional blocking solution (cBS)

cBS is composed of 2% BSA and 0.5% Normal Donkey Serum in PBS. BSA (Sigma-Aldrich), Normal Donkey Serum (Abcam).

### Imaging buffer (IB)

IB is composed of 700mM N-Acetyl-Cysteine (NAC) in ddH_2_O and adjusted to pH 7.4. NAC (Sigma-Aldrich).

### Elution buffer (EB)

EB is composed of 0.5M L-Glycine, 3M Guanidium chloride (GC), 3M Urea and 70mM Tris(2-carboxyethyl)phosphine hydrochloride (TCEP) in ddH_2_O adjusted to pH 2.5. L-Glycine (Sigma-Aldrich), GC (Sigma-Aldrich), Urea (Sigma-Aldrich), TCEP (Sigma-Aldrich).

### Primary antibodies

Antibodies used in this study were selected based on the following characteristics: (1) Antibody has been successfully used in immunofluorescence (IF) and published in scientific literature. (2) Changes in signal intensity, texture and localization in IF experiments upon treatment with EGF and/or pharmacological inhibition are coherent with current consensus beliefs. (3) Antibodies can successfully be eluted, and no residual signal is detected. Detailed information on antibodies can be found in Extended Data Table 1 and Extended Table 2.

### Secondary antibodies

Donkey-anti-Rabbit IgG Alexa Fluor 488, Donkey-anti-Mouse IgG Alexa Fluor 488, Donkey-anti-Rabbit IgG Alexa Fluor 568, Donkey-anti-Mouse IgG Alexa Fluor 568, were each diluted 1:500 in cBS. Donkey-anti-Rabbit IgG Alexa Fluor 488 (Thermo Fisher Scientific), Donkey-anti-Mouse IgG Alexa Fluor 488 (Thermo Fisher Scientific), Donkey-anti-Rabbit IgG Alexa Fluor 568 (Thermo Fisher Scientific), Donkey-anti-Mouse IgG Alexa Fluor 568 (Thermo Fisher Scientific).

### DNA stain solution (DSS)

75µg/ml 4’, 6-Diamidino-2-phenylindole (DAPI) PBS was used to counter-stain DNA for each 4i cycle. DAPI (Thermo Fisher Scientific).

### Whole protein stain solution (WPSS)

WPSS is composed of 0.166µg/ml Succinimidyl Ester Alexa Fluor 647 (SUCCS) in ddH_2_O containing 100mM NaHCO_3_ and 2.5mM Na_2_CO_3_. SUCCS (Thermo Fisher Scientific).

### Computational infrastructure

Image analysis was performed on a high-performance cluster provided by Science Cloud UZH using TissueMAPS (https://github.com/pelkmanslab/TissueMAPS) and CellProfiler. All other described computational methods were executed on Virtual Machines provided by Science Cloud UZH.

## Methods

### Cell culture

184A1 cells were cultured in CGM at 37°C, 95% Humidity and 5% CO_2_. For experiments, 4.000 cells were seeded per well in a 96-well plate with plastic bottom (Thermo Fisher Scientific) and grown for 4 days in CGM at 37°C, 95% Humidity and 5% CO_2_.

### Growth factor stimulation

Prior to specific treatments, cells were depleted of external growth factors (HS, EGF and Insulin) and complex, undefined protein mixtures (HS). To that end, cells were washed 10 times with PBS and CGM replaced with AM and incubated for 12 hours at 37°C, 95% Humidity and 5% CO_2_. Cells were then treated with varying concentrations of EGF (0, 6.25, 10, 25 and 100ng/ml) diluted in AM for 5 min.

### Iterative Indirect Immunofluorescence Imaging (4i)

Sample preparation was carried out as following. After treatment, cells were fixed with 4% Paraformaldehyde (Electron Microscopy Sciences) in PBS at room temperature for 20 min and afterwards washed 5 times with PBS. Cells were then permeabilized using 0.25% Triton X-100 in PBS at room temperature for 10 min followed by 5 times washing with PBS. Subsequently, each of the following steps was performed at room temperature in order of their description for each cycle of 4i if not indicated otherwise. (1) Antibody Elution. The sample was washed 4 times with ddH_2_O which was then aspirated to the lowest possible volume. The following step was then repeated 3 times. EB was added to the sample and incubated on a see-saw rocker (SSR) for 10 min at 14 oscillations per minute and aspiration to the lowest possible volume. After those 3 iterations the sample was washed 5 times with PBS. (2) Blocking. sBS was added to the sample, transferred to a SSR for 1 hour at 14 oscillations per minute and afterwards washed 6 times with PBS. (3) Primary antibody binding. The primary antibody was diluted in cBS and added to the sample. After incubation for 2 hours on a SSR at 14 oscillations per minute the sample was washed 4 times with PBS. (4) Secondary antibody binding. The secondary antibody was diluted in cBS and added to the sample. After incubation for 1 hours on a SSR at 14 oscillations per minute the sample was washed 6 times with PBS. (5) DNA staining. DSS was prepared and added to the sample. After incubation for 10 min on a SSR at 14 oscillations per minute the sample was washed 5 times with ddH_2_O. (5.5) Whole protein staining (This step is only performed in the last 4i acquisition cycle). WPSS was prepared and added to the sample. After incubation for 5 min on a SSR at 14 oscillations per minute the sample was washed 5 times with ddH_2_O. (6) Imaging. IB was added to the sample, the plate covered with aluminum foil and subjected to imaging. All liquid dispensing and washing steps described for 4i were performed using a Washer Dispenser EL406 (BioTek).

### Microscopy

Acquisition of microscopy images was performed using an automated spinning disk microscope from Yokogawa (CellVoyager 7000) with an enhanced CSU-W1 spinning disk (Microlens-enhanced dual Nipkow disk confocal scanner, wide view type) in combination with a 20x Nikon air objective of 0.75 NA, and andor cSMOS cameras. 10 z-planes with a 500nm z-spacing were acquired per site and the maximum intensity projection was calculated and used for subsequent image analysis. UV (406nm), green (488nm), red (568) and far-red (647nm) signals were acquired sequentially.

### Image alignment for acquisitions from different 4i cycles

Slight shifts in X and Y can occur between acquisition cycles due to imperfect stage repositioning. Therefore, microscopy images of different cycles from the same site require image alignment. DAPI images from across all cycles were used for image registration relative to cycle 01. The calculated shifts were then applied to acquisitions from 405nm, 488nm, 568nm and 647nm resulting in aligned imaging sites. The aligned images were then used for subsequent image processing, object identification and feature extraction. Image alignment was performed using TissueMAPS.

### Image processing, object identification and feature extraction

TissueMAPS and CellProfiler^67^ 3 (CP) were used to perform image processing, object identification and feature extraction with standard CP and custom TissueMAPS modules. Every image was corrected for illumination biases prior to alignment in TissueMAPS using summary statistics for each individual cycle and acquisition and the constant background substracted. Objects were then segmented using CP. Nuclei (primary object) were identified using images of DAPI. Further object segmentation was performed by combining signals from Calreticulin (cycle 12) and SUCCS (cycle 20) in a 1:1 ratio for the same site. Watershed algorithm was then used to identify the outline of Cells (secondary object). Further objects (tertiary objects) were derived from both primary and secondary objects. Cytoplasm is representing the region where the cell mask is present, but the nucleus mask is not. The membrane object is the difference between the full cell mask and a cell mask shrunken by 10 pixels. Cytobody object is the region where the cytoplasm mask is present, but the membrane mask is not. Label images for all objects were derived and used in TissueMAPS to extract features for all single cells. Intensity, texture and area shape of objects were extracted. Population features were calculated on label images for objects using custom MATLAB scripts.

### Classification of miss-segmented, mitotic, multinucleated cells (Quality control)

A single, two-class Support Vector Machine (SVM) classifier was employed to separate miss-segmented, mitotic and multinucleated cells from cells not exhibiting those characteristics. An initial training set was defined by manual assignment of representative cells to each class. Texture and intensity features of DAPI images and shape properties of the nucleus were used as independent variables. Iterative training and prediction accompanied with visual inspection was performed till sufficient accuracy was achieved. A final prediction step was then performed, and the outcome used to exclude miss-segmented, mitotic and multinucleated cells from further analysis. Cells touching an image border were identified by automated discretion and also excluded from analysis.

### Data analysis

Data analysis was performed with custom scripts in MATLAB (2017b), R (3.4) and Python (2.7) using inbuild functions and custom functions when indicated.

### Data transformation and normalization

Data derived from response marker staining panel were log-transformed (base 2) after image processing and the transformed values used for analysis except in Extended Data Fig. 1e and EC50 calculation where non-transformed values where used. Data derived from the cellular state marker staining were both used as untransformed and log-transformed (base 2) values after image processing. For Extended Data Fig. 1e the untransformed values were used. Antibody derived values for the cellular state (texture and intensity features) were z-scored per well for both transformed and untransformed values. Segmentation derived features (morphology and population features) were z-scored per plate. Of these, only morphology features were used both with their untransformed and log-transformed value (base 2).

### Kernel density estimation

Kernel density estimation was performed with inbuild functions of MATLAB using a gaussian kernel and a bandwidth parameter according to Silverman’s rule of thumb. When used, weights for density estimation are indicated.

### Calculation of the overlap coefficient between replicates

To assess the reproducibility of the independent replicates the shared overlap area between two distributions was calculated. To achieve this, an empirical probability density function was estimated using kernel density estimation. Monte-Carlo integration was then used over a grid to approximate the shared area under the empirical density distributions. Shared overlap area was then calculated as fraction of shared area against the joined individual areas of each distribution and is termed overlap coefficient. Calculation was performed on log_2_ transformed but non-normalized values after image processing.

### Merging of independent replicates for data analysis

Due to the high reproducibility of the independent replicates, single-cell data from replicates were merged for most analysis performed. For estimation of the coefficient of determination (R^2^) and prediction of classes, this generally yields worse estimates as no additional term was included to account for technical variability.

### Linear and logistic regression

Regression modeling was performed using the Glmnet package^68^. Cross-validation was performed during training 10 times. Regularization path was computed for the elastic-net penalty across a grid of different values for the regularization parameter lambda. For prediction the most regularized model (in terms of lambda) within one standard error of the minimum was used. When used for prediction, randomly subsampled cells were used for training and the remaining cells used for prediction. Bootstrapping was performed and is indicated whenever a prediction for every cell in the data-set was needed. Independent and dependent variables are indicated at the respective points.

### Significance tests

Whenever used, the performed statistical test, the null hypothesis and the significance thresholds are indicated in the figure legends.

### Calculation of unique and shared variance explained

The aim of this analysis is to quantify whether complementary different sets of independent variables explain the same or different variability observed in the dependent variable^69^. To achieve this linear regression was performed with a “complete” set of independent variables (i.e. the cellular state and other response markers as indicated) and the adjusted coefficient of determination (R^2^) calculated. Further models were trained with subsets of independent variables, namely the cellular state or other response markers alone and the R^2^ calculated. Unique variability explained by one specific subset is calculated as the R^2^ difference between the model on all independent variables and the model on one subset. This results in the fraction of variability which is additionally, and hence uniquely, explained by the other subset of independent variables. Shared variability explained is consequently the difference between highest achievable R^2^, and the combined unique variability explained by each subset. Increase in R^2^ due to interactions between the variable sets is here treated as a unique increase as the gain in variance explained is only possible when the other variable set is present.

### Correlation and partial correlation analysis

Linear correlation and partial correlation coefficients were calculated using inbuild functions of MATLAB (corr and partialcorr respectively). If not indicated otherwise, Pearson’s r was calculated.

### Non-redundant, unique association with specific cellular state features

To estimate the unique association of each signaling response marker with cellular state features, the feature set was first reduced to a readily interpretable subset (Extended Data Table 4). Then the partial correlations of each response marker with each of those features controlled for the remaining response markers was calculated.

### Similarity of association patterns across concentrations of EGF

To assess how similar the non-redundant association pattern of response markers with cellular state features is across concentrations, first the partial correlations for each response marker and each concentration was calculated as described. This yields a vector of partial correlation coefficients to cellular state features for each concentration. Then the cosine similarity between these vectors was calculated in a pairwise comparison for combination of concentrations.

### Estimation of cellular state similarity of neighbors in signaling space

For each cell in the data, separated by concentration of EGF, the closest 50 neighbors were calculated based either on the complete set of response markers or each individual response marker alone. Manhattan distance was used to calculate the closest neighbors in signaling space. From these data 20 cells were randomly sampled 10^5^ times and the mean cosine similarity in the cellular state of the cells compared to their neighbors calculated.

### Fuzzy clustering

Fuzzy clustering assigns observations non-exclusively to different clusters. It was here employed as the data are defined by both “discrete” and “continuous” properties. The inbuild function c-means of MATLAB was used with modifications to accept Manhattan distance as metric during clustering. Before clustering, a set of principal components (PC) which describe the cellular state space and are linked to the multimodal signaling response were identified using sequential feature elimination with regression analysis. PCs which were estimated to be significant predictors in 90% of all models were chosen for cluster analysis. Further, PCs were weighted based on the mean regression coefficients (more precisely, their mean absolute values) across all models to ensure clustering in relevant dimensions. The heuristic for an optimal number of centroids was here estimated using two separate methods. 1. Self-organized maps^70^ (SOM) were used for clustering using a different starting numbers and grids of neurons. The Bayesian Information Criterion was then calculated for each clustering results. This yielded 18 as a consensus optimum for the number of centroids. 2. PhenoGraph^71^ based clustering was employed across different numbers of neighbors to build the initial neighborhood graph. Subsequent community detection identified two plateaus at 20 and 15. In combination with the SOM results, 18 was therefore chosen as the number of centroids for clustering. Fuzzy clustering further requires a weighting exponent coefficient (WEC) which influences the centroid positions and the fuzziness vs. crispness of the resulting membership coefficients. The Gini-coefficient, which can be used to describe how unequal the membership coefficients are distributed for each cell, was calculated for a range of WECs. A value of 1.125 was identified as a consensus between an unequal value distribution of the membership coefficient yet sufficient fuzziness of the clustering.

### EC50 analysis

To estimate the cellular-state dependent sensitivities of each response marker, EC50 analysis was performed. To achieve this, the weighted mean response on non-transformed values of each cluster was calculated for each concentration of EGF. These values were normalized between 0-1 and a 4-parameter logistic regression against the concentration of EGF performed. The parameter which corresponds to the inflection point of the resulting S-shaped curve was extracted and represents the EC50.

## Data Availability

All raw and processed data can be provided upon reasonable request.

## Code Availability

All scripts used for data analysis can be provided upon reasonable request.

---

**Extended Data Table 1:**
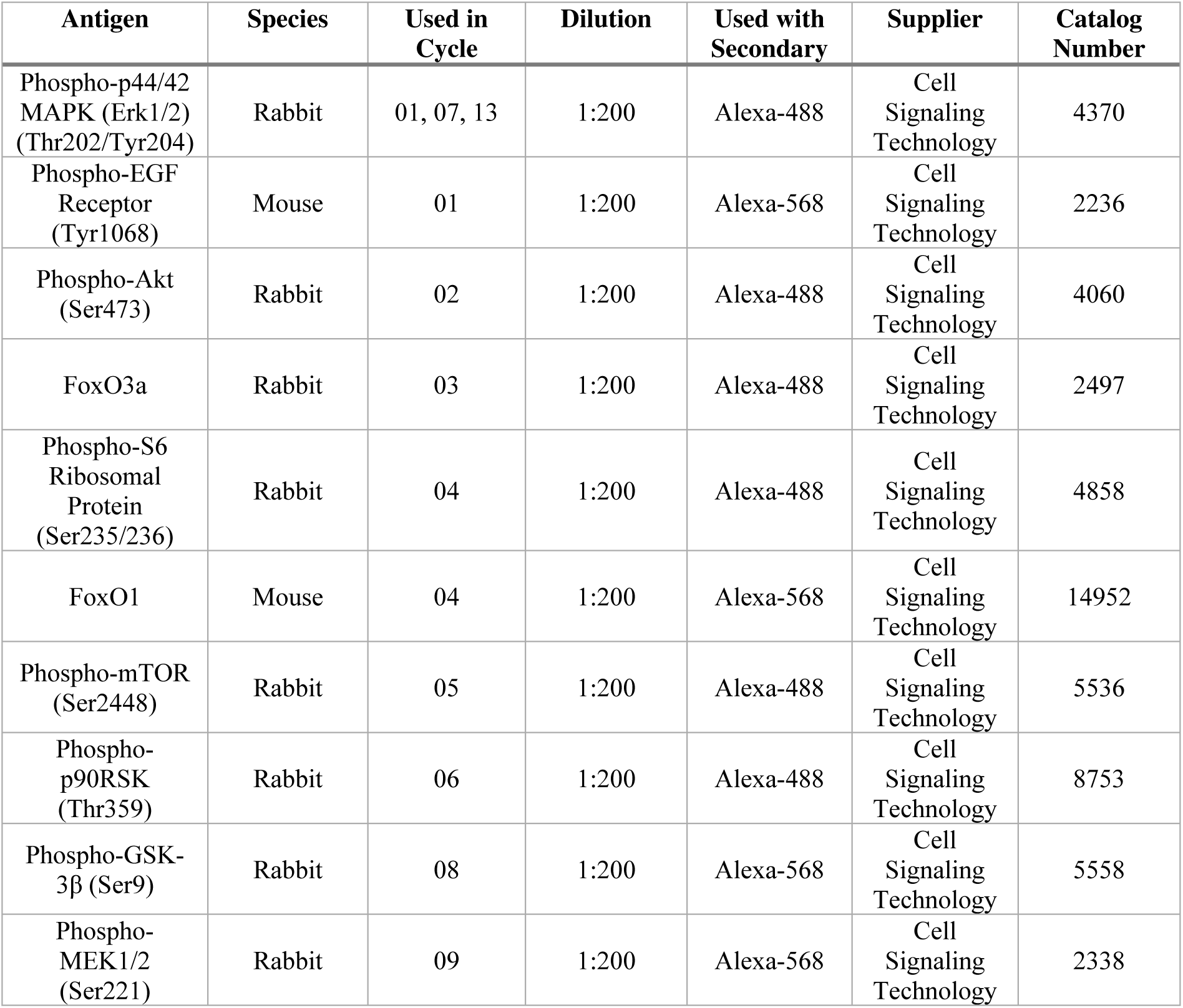
Signaling response marker.

| **Antigen** | **Species** | **Used in Cycle** | **Dilution** | **Used with Secondary** | **Supplier** | **Catalog Number** |
| --- | --- | --- | --- | --- | --- | --- |
| Phospho-p44/42 MAPK (Erk1/2) (Thr202/Tyr204) | Rabbit | 01, 07, 13 | 1:200 | Alexa-488 | Cell Signaling Technology | 4370 |
| Phospho-EGF Receptor (Tyr1068) | Mouse | 01 | 1:200 | Alexa-568 | Cell Signaling Technology | 2236 |
| Phospho-Akt (Ser473) | Rabbit | 02 | 1:200 | Alexa-488 | Cell Signaling Technology | 4060 |
| FoxO3a | Rabbit | 03 | 1:200 | Alexa-488 | Cell Signaling Technology | 2497 |
| Phospho-S6 Ribosomal Protein (Ser235/236) | Rabbit | 04 | 1:200 | Alexa-488 | Cell Signaling Technology | 4858 |
| FoxO1 | Mouse | 04 | 1:200 | Alexa-568 | Cell Signaling Technology | 14952 |
| Phospho-mTOR (Ser2448) | Rabbit | 05 | 1:200 | Alexa-488 | Cell Signaling Technology | 5536 |
| Phospho-p90RSK (Thr359) | Rabbit | 06 | 1:200 | Alexa-488 | Cell Signaling Technology | 8753 |
| Phospho-GSK-3β (Ser9) | Rabbit | 08 | 1:200 | Alexa-568 | Cell Signaling Technology | 5558 |
| Phospho-MEK1/2 (Ser221) | Rabbit | 09 | 1:200 | Alexa-568 | Cell Signaling Technology | 2338 |

**Extended Data Table 2:** Cellular state marker.

| **Antigen** | **Organelle/**<br>**Function** | **Species** | **Used**<br>**in**<br>**Cycle** | **Dilution** | **Used with**<br>**Secondary** | **Supplier** | **Catalog**<br>**Number** |
| --- | --- | --- | --- | --- | --- | --- | --- |
| Paxillin | Focal Adhesions | Mouse | 03 | 1:200 | Alexa-568 | Thermo-Fisher Scientific | AHO0492 |
| GM130 | Golgi | Mouse | 05 | 1:200 | Alexa-568 | BD Biosciences | 610823 |
| β-Actin | Cytoskeleton | Mouse | 08 | 1:400 | Alexa-488 | Abcam | ab8226 |
| EEA1 | Endosomes | Mouse | 09 | 1:200 | Alexa-488 | BD Biosciences | 610457 |
| ERGIC-53 | ER-Golgi | Mouse | 10 | 1:200 | Alexa-488 | Enzo Life Sciences | ENZ-ABS300-0100 |
| Calreticulin | ER | Rabbit | 12 | 1:200 | Alexa-568 | Abcam | ab2907 |
| ABCD3 | Peroxisomes | Mouse | 14 | 1:300 | Alexa-568 | Merck | AMAB90995 |
| Sec13 | Secretory Pathway | Rabbit | 14 | 1:500 | Alexa-488 | Lubio Science | A303-980A |
| Dynamin 2 | Endocytosis | Rabbit | 15 | 1:300 | Alexa-488 | Abcam | ab3457 |
| Yap1 | Mechanical State | Mouse | 15 | 1:400 | Alexa-568 | Santa Cruz Biotechnology | sc-101199 |
| HSP60 | Mitochondria | Mouse | 16 | 1:500 | Alexa-488 | Abcam | ab13532 |
| Cyclin B1 | Cell Cycle | Rabbit | 16 | 1:200 | Alexa-568 | Cell Signaling Technology | 12231 |
| PCNA | Cell Cycle | Rabbit | 17 | 1:200 | Alexa-488 | Cell Signaling Technology | 13110 |
| Polymerase II CTD repeat YSPTSPS (phospho S5) | Transcription | Mouse | 17 | 1:200 | Alexa-647 | Abcam | ab5408 |
| VPS35 | Endosomes | Goat | 19 | 1:400 | Alexa-488 | Novus Biologicals | IMG-3575 |
| Phospho-Rb (Ser807/811) | Proliferation | Rabbit | 19 | 1:200 | Alexa-568 | Cell Signaling Technology | 8516 |
| α-Tubulin | Cytoskeleton | Mouse | 20 | 1:500 | Alexa-488 | Abcam | ab7291 |
| DDX6 | P-Bodies | Rabbit | 20 | 1:500 | Alexa-568 | Lubio Science | A300-461A |

**Extended Data Table 3:** Intensity and object assessed.

| Marker | Object quantified | Intensity type |
| --- | --- | --- |
| **Signaling response** |  |  |
| pEGFR | Cytoplasm | Mean intensity |
| pMEK | Cytoplasm | Mean intensity |
| pERK | Cytoplasm | Mean intensity |
| pRSK | Cytoplasm | Mean intensity |
| pGSK3B | Cytoplasm | Mean intensity |
| pMTOR | Cytoplasm | Mean intensity |
| pAKT | Cytoplasm | Mean intensity |
| FoxO1 | Nucleus | Mean intensity |
| FoxO3a | Nucleus | Mean intensity |
| pS6 | Cytoplasm | Mean intensity |
| **Cellular state** |  |  |
| Paxillin | Cytoplasm | Mean intensity |
| GM130 | Cytoplasm | Mean intensity |
| β-Actin | Cytoplasm | Mean intensity |
| EEA1 | Cytoplasm | Mean intensity |
| ERGIC-53 | Cytoplasm | Mean intensity |
| Calreticulin | Cytoplasm | Mean intensity |
| ABCD3 | Cytoplasm | Mean intensity |
| Sec13 | Cytoplasm | Mean intensity |
| Dynamin | Cytoplasm | Mean intensity |
| YAP | Nucleus | Mean intensity |
| HSP60 | Cytoplasm | Mean intensity |
| CyclinB | Cytoplasm | Mean intensity |
| PCNA | Nucleus | Mean intensity |
| pPOLII | Nucleus | Mean intensity |
| VPS35 | Cytoplasm | Mean intensity |
| pRB | Cytoplasm | Mean intensity |
| Tubulin | Cytoplasm | Mean intensity |
| DDX6 | Cytoplasm | Mean intensity |

**Extended Data Table 4:** Subset of cellular state features.

| Feature Type | Name | Feature description |
| --- | --- | --- |
| **Population Features** | Cell contact | Cell contact - Percentage of membrane in contact with other cells |
|  | LCD1 | Local cell density 1 – Density of other nuclei in proximity |
|  | LCD2 | Local cell density 2– Density of other nuclei in an extended region |
|  | PCD | Population density – Density of other cell bodies in proximity |
| **Morphology Features** | Area (Nucleus) | Log<sub>2</sub> Nuclear Area – Area occupied by a cell's nucleus |
|  | Area (Cell) | Log<sub>2</sub> Cell Area – Area occupied by a cell |
|  | Perimeter (Cell) | Log<sub>2</sub> Cell Perimeter – Amount of membrane |
|  | Roundness (Cell) | Log<sub>2</sub> Cell Roundness – Indicator describing the roundness of a cell |
| **Intensity Features** | DAPI | Log<sub>2</sub> Nucleus mean DAPI – Concentration of DNA in a cell's nucleus |
|  | Paxillin | Log<sub>2</sub> Cytoplasm mean Paxillin – Abundance (Concentration) of Paxillin in a cell's cytoplasm |
|  | GM130 | Log<sub>2</sub> Cytoplasm mean GM130 – Abundance (Concentration) of the Golgi in a cell's cytoplasm |
| | Actin | Log<sub>2</sub> Cytoplasm mean Actin – Abundance (Concentration) of α-Actin in a cell's cytoplasm |
|  | EEA1 | Log<sub>2</sub> Cytoplasm mean EEA1 – Abundance (Concentration) of early endosomes in a cell's cytoplasm |
|  | ERGIC53 | Log<sub>2</sub> Cytoplasm mean ERGIC-53 – Abundance (Concentration) of the ERGIC in a cell's cytoplasm |
|  | Calreticulin | Log<sub>2</sub> Cytoplasm mean Calreticulin – Abundance (Concentration) of ER in a cell's cytoplasm |
|  | Sec13 | Log<sub>2</sub> Cytoplasm mean Sec13 – Abundance (Concentration) of a secretory pathway component in a cell's cytoplasm |
|  | ABCD3 | Log<sub>2</sub> Cytoplasm mean ABCD3 – Abundance (Concentration) of peroxisomes in a cell's cytoplasm |
|  | Dynamin | Log<sub>2</sub> Cytoplasm mean Dynamin – Abundance (Concentration) of Dynamin in a cell's cytoplasm |
|  | YAP | Log<sub>2</sub> Nucleus mean YAP – Abundance (Concentration) of YAP in a cell's nucleus |
|  | HSP60 | Log<sub>2</sub> Cytoplasm mean HSP60 – Abundance (Concentration) of mitochondria in a cell's cytoplasm |
|  | CyclinB | Log<sub>2</sub> Cytoplasm mean CyclinB – Abundance (Concentration) of CyclinB in a cell's cytoplasm |
|  | PCNA | Log<sub>2</sub> Nucleus mean PCNA – Abundance (Concentration) of PCNA in a cell's nucleus |
|  | pPOLII | Log<sub>2</sub> Nucleus mean pPOLII – Abundance (Concentration) of pPOLII in a cell's nucleus |
|  | VPS35 | Log<sub>2</sub> Cytoplasm mean VPS35 – Abundance (Concentration) of late endosomes in a cell's cytoplasm |
|  | pRB | Log<sub>2</sub> Nucleus mean pRB – Abundance (Concentration) of pRB in a cell's nucleus |
|  | Tubulin | Log<sub>2</sub> Cytoplasm mean Tubulin – Abundance (Concentration) of microtubules in a cell's cytoplasm |
|  | DDX6 | Log<sub>2</sub> Cytoplasm mean DDX6 – Abundance (Concentration) of P-Bodies |

